# Endothelial YAP/TAZ rewiring under cardiometabolic stress drives sex-divergent vascular remodeling in heart failure with preserved ejection fraction

**DOI:** 10.64898/2026.05.07.723127

**Authors:** Alexandra Klaus-Bergmann, Laura Katharina Sievers, Jakob Versnjak, Katharina Koch, Tomasz Nawara, Eireen Bartels-Klein, Oliver Popp, January Weiner, Katja Meier, Irene Hollfinger, Ilona Kamer, Martin Taube, Arnd Heuser, Tatiana Borodina, Dieter Beule, Michael Potente, Ulf Landmesser, Philipp Mertins, Marcus Kelm, Dominik N. Müller, Holger Gerhardt

**Affiliations:** Integrative Vascular Biology Lab, Max Delbrück Center for Molecular Medicine in the Helmholtz Association (MDC), Berlin, Germany; Experimental and Clinical Research Center, a cooperation of Charité-Universitätsmedizin Berlin and Max Delbrück Center for Molecular Medicine, Berlin, Germany; Medical Department IV-Nephrology and Hypertension, UKSH, Kiel, Germany; Institute of Computer-assisted Cardiovascular Medicine, Deutsches Herzzentrum der Charité, Berlin, Germany; Charité-Universitätsmedizin Berlin, corporate member of Freie Universität Berlin and Humboldt-Universität zu Berlin, Berlin, Germany; Proteomics Platform, Max-Delbrück-Center for Molecular Medicine in the Helmholtz Association (MDC), Berlin, Germany; Proteomics Platform, Berlin Institute of Health (BIH), Charité – Universitätsmedizin Berlin, Berlin, Germany; Genomics Technology Platform, Max-Delbrück-Center for Molecular Medicine in the Helmholtz Association (MDC), Berlin Institute for Medical Systems Biology (BIMSB), 10115 Berlin, Germany; Translational Bioinformatics, Max Delbrück Center for Molecular Medicine in the Helmholtz Association (MDC), Berlin, Germany; Animal Phenotyping, Max Delbrück Center for Molecular Medicine in the Helmholtz Association (MDC), Berlin, Germany; Deutsches Herzzentrum der Charité, Department of Cardiology, Angiology and Intensive Care Medicine, Campus Benjamin Franklin, Berlin, Germany; Angiogenesis & Metabolism Laboratory, Max Delbrück Center for Molecular Medicine in the Helmholtz Association (MDC), Berlin, Germany; Department of Congenital Heart Disease – Pediatric Cardiology, Deutsches Herzzentrum der Charité, Berlin, Germany; DZHK (German Centre for Cardiovascular Research), partner site Berlin, Berlin, Germany; Helmholtz Institute for Translational AngioCardiosciences (HI-TAC), Max Delbrück Center for Molecular Medicine at Heidelberg University, Heidelberg, Germany; Friede Springer Cardiovascular Prevention Center at Charité - Berlin, Germany; Berlin Institute of Health at Charité (BIH), Center of Vascular Biomedicine, Universitätsmedizin Berlin, Berlin, Germany

**Keywords:** Heart failure, endothelial dysfunction, mechanobiology, vascular remodelling, plasma proteomics

## Abstract

Heart failure with preserved ejection fraction (HFpEF) is widely linked to endothelial dysfunction, yet the molecular pathways translating cardiometabolic stress into microvascular remodeling remain poorly defined. Here, we identify endothelial YAP/TAZ signaling as a mechanistic regulator of sex-divergent vascular responses in HFpEF. Plasma proteomics from the UK Biobank revealed elevated circulating YAP1 levels associated with heart failure and increased mortality, particularly in male patients, where YAP1 coincided with increased levels of the endothelial activation marker ESM1. In a hypertensive cardiorenal mouse model, endothelial YAP/TAZ deletion preserved cardiac function, whereas endothelial TAZ gain-of-function aggravated disease. Under cardiometabolic stress (TNFα and high glucose), endothelial cells exhibited sex-specific rewiring of YAP/TAZ-dependent transcriptional programs. Male endothelial cells showed increased extracellular YAP1 release, angiogenic instability with impaired extracellular matrix remodeling, whereas female cells adopted an immune-primed, stress-adaptive phenotype. Mechanistically, cardiometabolic stress uncoupled canonical YAP-TEAD transcription and engaged alternative cofactors, including VGLL3 and VGLL4, thereby reshaping the endothelial secretome and propagating sex-divergent microvascular remodeling. These findings identify endothelial YAP/TAZ rewiring as a molecular switch that converts cardiometabolic stress into sex-divergent microvascular remodeling in HFpEF and connect this process to circulating YAP1 and ESM1 in patients.

## Introduction

HFpEF is an increasingly prevalent clinical syndrome, accounting for the majority of heart failure cases. Despite the growing burden, therapeutic progress remains limited, largely due to the heterogeneity of its pathophysiology and the lack of effective stratification tools^1^. Increasing evidence identifies systemic endothelial dysfunction as a central feature of HFpEF, closely associated with comorbidities such as hypertension, obesity, and diabetes. However, the molecular pathways through which these stressors disrupt endothelial function and contribute to myocardial remodeling remain poorly defined.

The endothelium is not a passive bystander but an active transducer of biomechanical and metabolic cues. Among the signaling pathways that integrate these stress inputs, the transcriptional co-activators YAP1 and WWRT1/TAZ (hereafter referred to as YAP/TAZ) have emerged as key regulators of endothelial gene expression programs controlling proliferation, survival, differentiation, and migration. Through context-dependent nuclear translocation and interaction with transcription factors such as TEAD, YAP/TAZ coordinate tissue growth, endothelial morphology, vascular function, and adaptive cellular endocrine and paracrine responses to microenvironmental changes^2–10^. YAP/TAZ signalling has emerged as an important regulator of cardiac development and disease and is increasingly considered a potential therapeutic target for cardiac disease (recently reviewed^11^). However, most current knowledge derives from studies of cardiomyocytes^11,12^, epicardial cells^13^, and immune cells^14,15^. Despite evidence that endothelial YAP/TAZ regulates vascular inflammation, endothelial turnover, and vessel homeostasis, its role remains poorly understood and appears highly context-dependent^11,16,17^. Aberrant endothelial YAP/TAZ signaling has been implicated in atherosclerosis, chronic course of stroke, and vascular remodeling^17–20^; however, a specific role in HFpEF has not been defined. Current concepts of endothelial dysfunction in HFpEF assume that paracrine effects of the endothelium shape the perivascular microenvironment, driving inflammation, fibrosis, and myocardial stiffening^21–24^. The detailed molecular mechanisms that drive endothelial dysfunction and propagate endothelial stress to the perivascular niche remain poorly defined.

Here, we identify endothelial YAP/TAZ signaling as both a driver and biomarker of HFpEF. We identify elevated plasma YAP1 levels in human heart failure, which correlate with mortality. Using endothelial-specific mouse models, we show that loss of YAP/TAZ confers protection against heart failure development, while gain-of-function exacerbates disease, especially in male animals. Mechanistically, distinct cardiometabolic stressors (glycolytic constraint, or glucose excess and inflammation) induce a transcriptional co-activator switch in ECs, with sex-specific differences in downstream signaling, vascular integrity, and paracrine effects. Together, our data identify endothelial YAP/TAZ rewiring under cardiometabolic stress as a mechanistic determinant of sex-divergent vascular remodeling in HFpEF.

## Results

### Circulating Yap1 and endothelial markers are elevated in HFpEF patients

To explore circulating signatures associated with endothelial stress in heart failure (HF), we analyzed plasma proteomic profiles from the UK Biobank cohort^25^. Unexpectedly, circulating levels of the mechanotransducer YAP1 were significantly elevated in patients with heart failure with preserved ejection fraction (HFpEF) (Fig. 1A). Elevated YAP1 levels were accompanied by increased levels of several endothelial-associated proteins, including ESM1, CDH5, EDN1, ANGPT2, and VCAM1, consistent with endothelial activation. Comparison of plasma YAP1 levels across patient groups with obesity, diabetes, HFpEF, arterial hypertension (aHTN), and HF with reduced ejection fraction (HFrEF) revealed lower YAP1 levels in obese and diabetic individuals, whereas elevated levels were observed in HFpEF patients.

**Figure 1.**
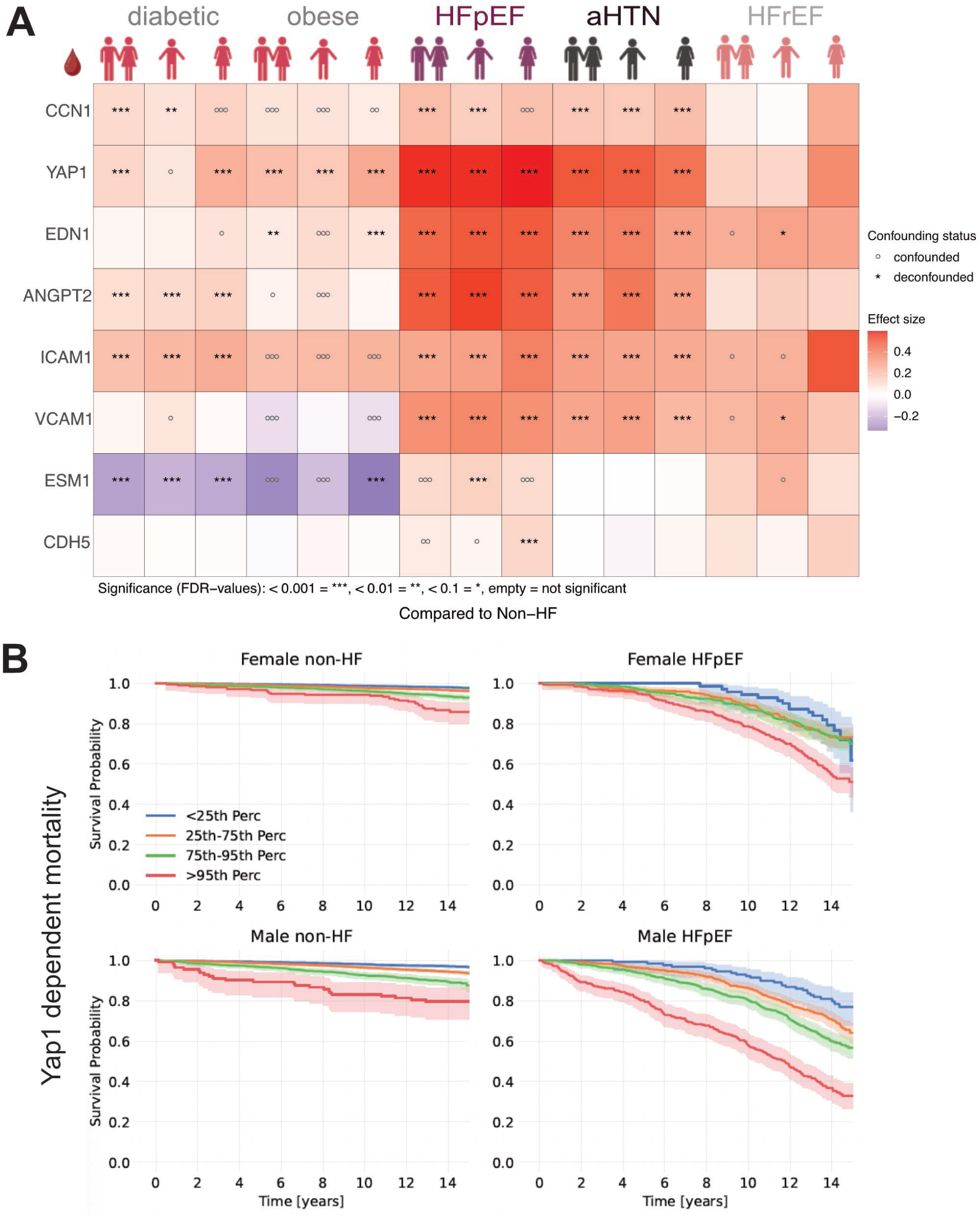
Circulating YAP1 is elevated in HFpEF and predicts mortality, particularly in men. (**A**) Heatmap showing the association of YAP1-TEAD, endothelial and HF-associated proteins in plasma of UKB patients using Olink3000 and was tested for potential confounding variables: circles (o) denote confounding features and asterisks (*) indicate deconfounded features. Statistical significance is indicated as follows: false discovery rate (FDR): <0.1 (*), <0.01 (**), <0.001 (***), empty cells represent FDR ≥0.1. (**B**) Kaplan-Meier survival curves stratified by YAP1 expression levels and clinical subgroups in men and women. Each curve represents one of four YAP1 expression ranges: below the 25_th_ percentile (blue), within the interquartile range (25_th_-75_th_ percentile, orange), between the 75_th_ and 95_th_ percentile, and above the 95_th_ percentile. Columns correspond to clinical groups: non-HF and HFpEF. Rows represent stratification by sex (female on top, male on bottom). Shaded regions indicate 95% confidence intervals.

Sex-stratified analysis uncovered divergent regulation of endothelial-derived plasma proteins in HFpEF, including ESM1, a marker associated with sprouting and angiogenic endothelium^26–28^, and CDH5 (VE-cadherin), a key component of endothelial junctional integrity^3,29,30^. In male patients with HFpEF, ESM1 levels were markedly elevated, but were attenuated in individuals with concomitant hypertension (Suppl. Fig.S1A). In contrast, female HFpEF patients exhibited significantly increased CDH5 levels and the highest circulating YAP1 concentrations. ESM1 levels were reduced in obesity and diabetes alone.

Stratification of HFpEF patients according to circulating YAP1 levels (below the 25^th^ percentile, 25^th^–75^th^ percentile, 75^th^–95^th^ percentile, and above the 95^th^ percentile) revealed a strong association between elevated YAP1 levels and mortality. Among patients above the 95^th^ percentile, a cumultative all-cause mortality reached approximately 20% after five years in the unadjusted Kaplan-Meier analysis in males, whereas a comparable mortality rate was reached only after approximately nine years in female HFpEF patients with aHTN (Fig. 1B, Extended Data Fig. 1B). Male HFpEF patients without aHTN exhibited the highest mortality risk, reaching approximately 20% mortality after two years.

Together, these analyses identify circulating YAP1 and ESM1 as candidate plasma biomarkers associated with an endothelial stress signature and adverse outcomes in participants classified as HFpEF.

To determine whether endothelial YAP/TAZ signaling contributes causally to cardiac remodeling in hypertensive heart failure, we next examined the effects of endothelial YAP/TAZ loss- and gain-of-function in mice.

### Endothelial YAP/TAZ regulates cardiac remodeling and survival in hypertensive heart failure

Inducible endothelial-specific YAP/TAZ loss-of-function (YT-iECKO) mice and endothelial TAZ gain-of-function mice (T-iECGOF) were subjected to a model of hypertensive cardiorenal injury involving unilateral nephrectomy (UNX), angiotensin II (AngII) infusion, and high dietary salt intake (Fig. 2A)^31,32^.

**Figure 2.**
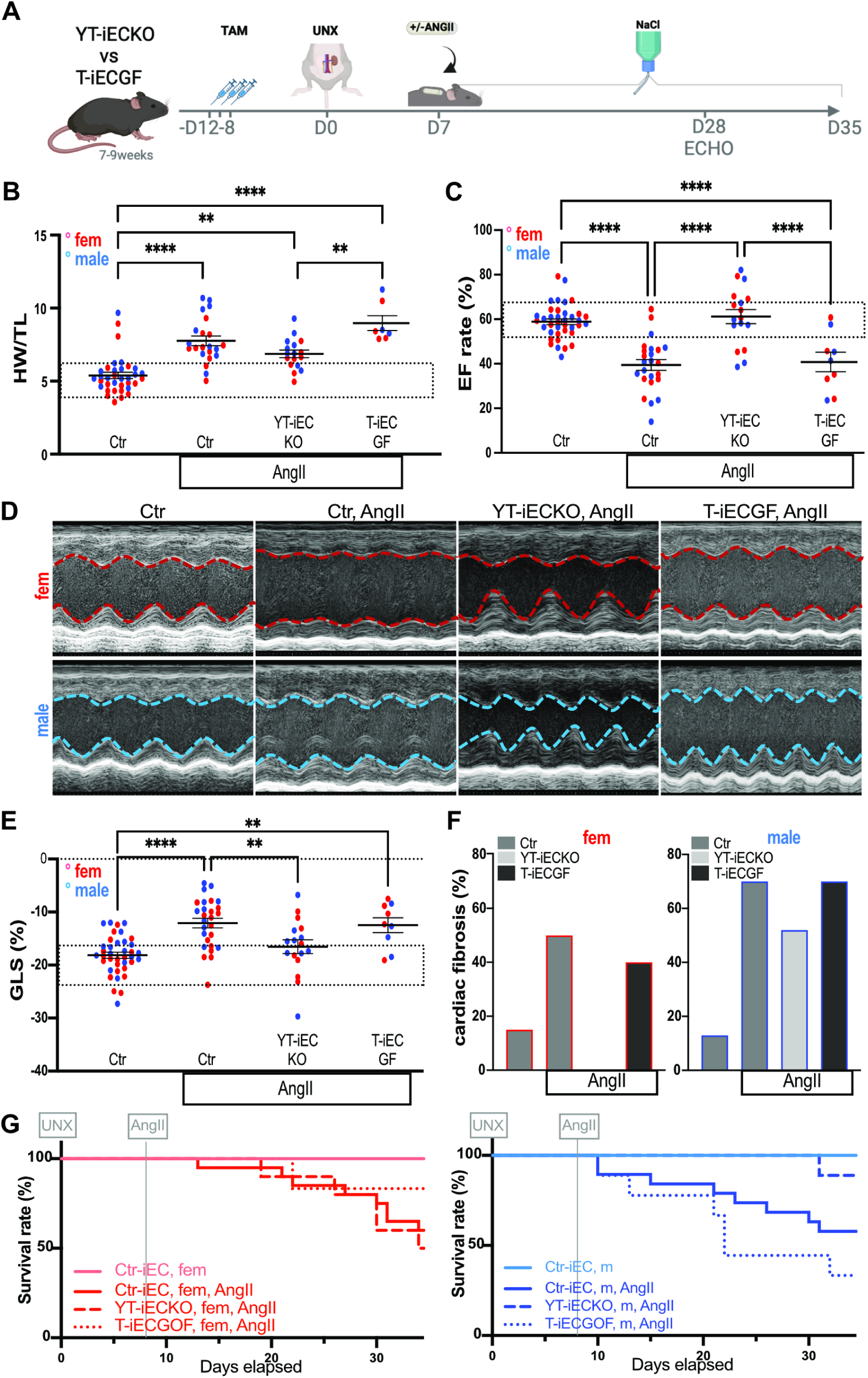
Endothelial YAP/TAZ loss is protective, while activation exacerbates HFpEF in vivo. (**A**) Schematic representation of the hypertensive cardiomyopathy and nephropathy mouse model (uninephrectomy (UNX) with or without Angiotensin II (AngII) pump implantation under the skin and salty (NaCl) water) was used for endothelial knock-out or gain of function of Yap or Taz (YT-iECKO, T-iECGF), created with. Therefore, mice received tamoxifen (TAM) 12, 10 and 8 days before UNX or sham surgery. (**B**) Tibia-length gross normalised heart weight (HW/TL). (**C-F**) Echocardiographic measures at 12-14 weeks: LV ejection fraction (EF-rate) (**C**), representative pictures of LV M-Mode (D), LV global longitudinal strain (GLS) (**E**) and quantification of cardiac fibrosis (blue, male; red, female) (F). (**G**) survival curves show increasing death in male T-iECGF and best survival in male YT-iECKO mice. Data displayed as mean ± SEM. One-way analysis of variance (ANOVA) followed by multiple-comparisons test. P values < 0.05 (*), <0.005 (**), <0.0005 (***), <0.0001 (****).

Following induction of hypertension, all experimental groups developed cardiac hypertrophy, as evidenced by increased heart weight-to-tibia length ratios (Fig. 2B). This hypertrophic response was accompanied by slightly reduced ejection fraction (EF; Fig. 2C), adverse left ventricular (LV) remodeling (Fig. 2D), and a deterioration of global longitudinal strain (GLS; Fig. 2E) in control and T-iECGOF mice. In contrast, these functional and structural impairments were largely absent in YT-iECKO animals.

Histological analyses revealed that AngII-induced epicardial and diffuse cardiac fibrosis was significantly reduced in female YT-iECKO mice (Fig. 2F, Extended Data Fig. 2A,B). Despite this reduction in fibrosis, approximately 50% of female mice subjected to the hypertensive cardiomyopathy and nephropathy protocol did not reach the predefined experimental endpoint but either died prematurely or required euthanasia due to welfare concerns. In contrast, male YT-iECKO mice showed markedly improved survival and were largely protected from AngII-induced excess mortality, despite the presence of myocardial fibrosis (Fig. 2G).

Together these results indicate that endothelial YAP/TAZ influences cardiac remodeling and survival under hypertensive stress, with distinct outcomes observed in female and male mice. To gain mechanistic insight into how cardiometabolic stress modulates endothelial YAP/TAZ signaling in a sex-specific manner, we next examined signaling responses in cultured endothelial cells.

### Cardiometabolic stress rewires YAP/TAZ signaling in female and male endothelial cells

Cardiometabolic stress responses were examined in pooled female and male human umbilical vein endothelial cells (HUVECs). Two complementary stress paradigms were applied: (i) glycolytic constraint using 2-deoxyglucose (2DG) to mimic metabolic energy restriction associated with ischemic stress, and (ii) combined TNFα and excess glucose (TNF/Glc) to model metabolic inflammation observed in cardiometabolic HFpEF (Fig. 3A–K).

**Figure 3.**
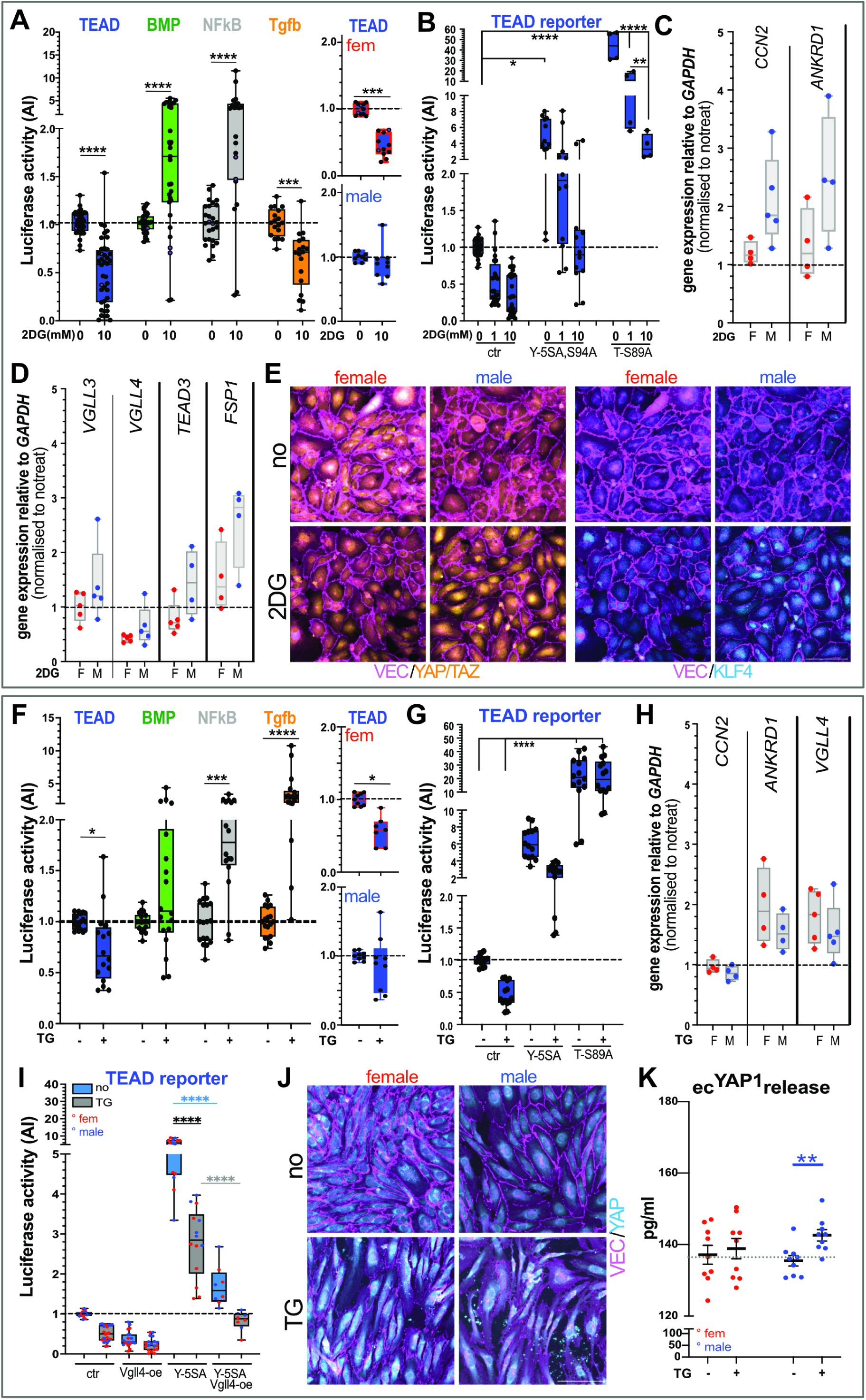
Cardiometabolic stress uncouples canonical YAP/TAZ–TEAD signaling and induces sex-specific rewiring. (**A-E**) Effect of glycolytic constraint on female and male HUVECs using 2-desoxyglucose (2DG). (**A,B)** Endothelial signaling and (**C,D)** gene response under glycolytic constraint shows female-specific decrease in YAP- and TAZ-dependent TEAD signaling while YAP/TAZ-TEAD target genes are normally expressed, suggesting shifts in the transcriptional TEAD response. In contrast, male HUVECs demonstrate elevated canonical TEAD target genes (**C**), *VGLL3*, *TEAD3* and *FSP1* expression pointing towards increased EndMT under 2DG (**D**). (**E**) Immunofluorescence analysis shows increased nuclear localization of YAP/TAZ (yellow; VE-Cadherin (VEC), magenta) in male and females under 2DG and a male-specific increase in nuclear KLF4 expression (cyan). Scale bar, 100µm. (**F-K**) Effect of cardiometabolic stress on female and male HUVECs using TNF/Glc (TG). (**F,G**) Endothelial signaling and (**H**) gene response under TNF/Glc shows female-specific decrease in the YAP-TEAD signaling response together with distinct expression of TEAD-dependent targets (**H**), suggesting shifts in YAP-dependent signaling. (**I**) Luciferase reporter assay for TEAD and the effect on YAP-TEAD signaling by VGLL4-overexpression (Vgll4-oe) under TNFGlc. (**J**) Immunofluorescence analysis showing nuclear localization of YAP (cyan) in female-ECs, but accumulation of speckle-like YAP-expression (scale bar, 100µm) and an increased release of YAP1 from male-ECs (ELISA, K). F/fem, female; M, male. Data displayed as mean ± SEM. Two-sided Wilcoxon t-test (paired, nonparametric) (A, F), one-way analysis of variance (ANOVA) followed by multiple-comparisons test (B-D, G-I). Mann-Whitney t-test (paired, non-parametric) (K). P values < 0.05 (*), <0.005 (**), <0.0005 (***), <0.0001 (****).

Under glycolytic constraint, luciferase reporter assays revealed a significant activation of BMP and NFκB signaling, accompanied by marked suppression of YAP/TAZ-TEAD and TGFβ reporter activity. Notably, suppression of TEAD reporter activity occurred selectively in female endothelial cells following 2DG treatment (Fig. 3A). Consistent with this observation, 2DG inhibited nuclear YAP/TAZ transcriptional activity by blocking constitutively active Yap-5SA/94SA and Taz-S89A in a dose-dependent manner (Fig. 3B). Despite reduced TEAD reporter activity, expression of canonical YAP/TAZ-TEAD target genes (e.g. *CCN2* and *ANKRD1*) remained detectable in female HUVECs and even increased in male cells (Fig. 3C). Male ECs showed increased expression of the TEAD-associated cofactor *VGLL3* together with *TEAD3* and *FSP1*, markers linked to endothelial-to-mesenchymal transition (EndMT)-like states^33–35^ (Fig. 3D). Glycolytic constraint also induced nuclear localization of YAP/TAZ in both sexes; however, increased nuclear accumulation of KLF4, a transcription factor downstream of BMP and shear stress signaling^36,37^, was observed only in male cells (Fig. 3E).

Exposure to TNF/Glc produced overlapping but distinct signaling responses. TEAD reporter activity was again selectively reduced in female endothelial cells, whereas both sexes showed robust activation of TGFβ signaling (Fig. 3F,G). Nuclear YAP-5SA-TEAD activity was suppressed, whereas nuclear activity of TAZ-S89A remained largely unaffected. Expression of a subset of YAP/TAZ-TEAD target genes including *ANKRD1*, increased following TNF/Glc treatment, whereas others such as *CCN2* remained unchanged, consistent with regulation by additional signaling pathways including TGFß or NFkB^38^. TNF/Glc treatment induced expression of *VGLL4*, a competitive inhibitor of YAP-TEAD interactions^39,40^ (Fig. 3H). Overexpression of VGLL4 effectively suppressed YAP-5SA-driven TEAD reporter activity under TNF/Glc conditions (Fig. 3I). TNF/Glc treatment also resulted in the appearance of YAP-positive extracellular speckle-like structures and was associated with increased YAP1 release from male endothelial cells (Fig. 3J,K).

Together, these experiments demonstrate that cardiometabolic stress alters endothelial YAP/TAZ signaling in a sex-dependent manner. Glycolytic stress preferentially suppressed canonical YAP/TAZ-TEAD activity in female endothelial cells, whereas male cells showed increased expression of alternative TEAD-associated factors including *VGLL3*, *TEAD3*, and KLF4. Under metabolic inflammatory stress, TNF/Glc reduced YAP-TEAD reporter activity while inducing VGLL4 expression and extracellular YAP1 release, particularly in male ECs. The release of YAP1 from stressed endothelial cells provides a potential cellular source for the elevated circulating YAP1 observed in HFpEF patients.

To define how these signaling alterations translate into broader endothelial stress programs, we next performed transcriptomic and proteomic analyses of endothelial cells exposed to TNF/Glc stress.

### TNF/Glc induces inflammatory activation with sex-specific endothelial programs

Bulk RNA sequencing and mass spectrometry–based proteomic profiling were performed in pooled female and male endothelial cells exposed to TNF/Glc. Under TNF/Glc exposure, ECs of both sexes exhibited a robust inflammatory core response. Transcriptomic analysis revealed activation of immune response pathways, cytokine signaling, interferon-stimulated gene programs, NFκB signaling, angiogenesis and transdifferentiation processes (Fig. 4). However, the relative balance between inflammatory activation, angiogenic remodeling, and cellular stress adaptation differed between sexes.

**Figure 4.**
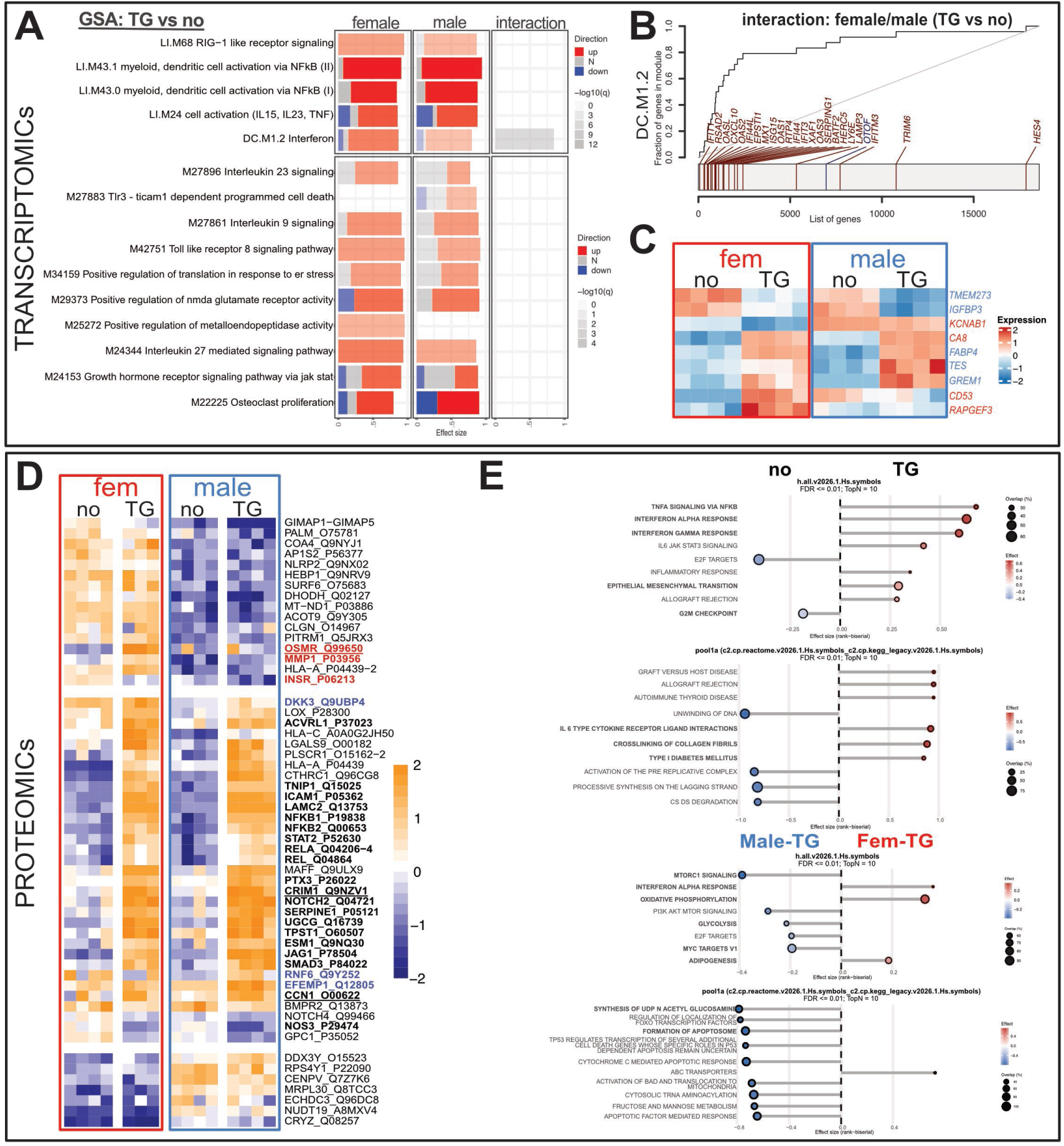
Male ECs adopt an unstable stress-driven angiogenic phenotype while female ECs activate an immune-primed but stress-adaptive state under HFpEF stimuli. (**A-C**) Bulk RNAsequencing results in female and male HUVECs under TNF/Glc (TG) show GSA results for the contrast TG vs no in female and male ECs and the interaction (fem_no_TG-fem_no_no)-(male_no_TG-male_no_no). Row corresponds to enriched molecular signature database term (tmod, hallmark, kegg, go), bar length indicates effect size (area under curve >0.8), intensity of the color corresponds to the FDR; red and blue fragments indicate fraction of DEGs within the contrast (FDR<0.1) (**A**). (**B**) Evidence plots of GO terms representing sex-differential regulation in interferon stimulated gene-modules (tmod ID DC.M1.2). X-axis is the list of all genes sorted by their p-value. Y-axis is the cumulative fraction. Light blue and light red colours represent significant down- or up-regulation. (**C**) Heatmaps display selected differentially expressed genes (log2FC > 0.4; FDR < 0.05). (**D-E**) Proteomic analysis of female and male ECs under TNF/Glc (TG) treatment. Heatmaps display selected differentially abundant proteins (log2FC > 1; adj. P < 0.05) (red, female samples; blue, male samples) (**D**). (**E**) Dot plot of significantly enriched hallmark, reactome, kegg pathways based on log2 fold changes (no vs TG and fem-TG vs. male-TG; n=4 no, n=3-4 TG). Enrichment was assessed using a two-sided Wilcoxon rank-sum test with Benjamini–Hochberg correction; pathways with FDR ≤ 0.05 are shown. The x-axis indicates the rank-biserial effect size (positive = higher in WT, negative = higher in dTGR). Point size represents the percentage of quantified proteins overlapping each pathway.

Gene set enrichment analysis indicated preferential activation of immune-associated pathways in female endothelial cells, including interferon-stimulated gene modules (tmod ID DC.M1.2), and cytokine signaling axes mediated by IL-23–*JAK2*, IL-9–*STAT1*, and IL-27–*OAS* (MSigDB IDs: M27896, M27861, M24344) (Fig. 4A,B). Consistent with this pattern, female ECs showed stronger induction of genes that support barrier function and adaptive remodelling^41,42^ including *RAPGEF3, CA8, CD53* and heightened suppression of *KCNAB1* following TNF/Glc exposure (Fig.4C). In contrast, male ECs exhibit gene expression changes associated with metabolic stress, lipid dysregulation and structural remodelling^43,44^ including pronounced upregulation in *FABP4, TES* and *GREM1* and alongside stronger downregulation in *IGFBP3* and *TMEM273*.

Proteomic profiling confirmed both shared and sex-divergent responses. In both sexes, TNF/Glc induced components of TGFβ-signaling (ACVRL1, SMAD3, SERPINE1), and NFκB signaling (NFκB1, NFkB2, REL, RELA), together with increased expression of endothelial activation markers including CCN1, CRIM1, ICAM1, ESM1, and PTX3 (Fig. 4D). 1D enrichment analysis similarly showed enrichment of inflammatory and cytokine signaling pathways, epithelial mesenchymal transition (EMT) and crosslinking of collagen fibrils in both sexes under TNF/Glc (Fig. 4E upper dot plot panels).

Beyond this shared inflammatory response, distinct sex-specific proteomic changes emerged. Female ECs showed increased OSMR and MMP1 expression together with reduced INSR levels. In contrast, male ECs preferentially induced RNF6 and EFEMP1, while these proteins displayed higher baseline expression in females (Fig. 4D). 1D enrichment analysis further revealed divergent pathway activation. Female ECs preferentially enriched pathways associated with interferon regulation, oxidative phosphorylation and adipogenesis, whereas male ECs showed enrichment of MYC target regulation, MTORC1 and glycolytic regulation, as well as apoptotic pathway regulation and posttranslational protein modification processes, including N-linked glycosylation (Fig. 4E, lower dot plot panels).

Together, these data indicate that TNF/Glc induces a shared inflammatory endothelial response while simultaneously engaging distinct sex-specific adaptive programs. Female ECs predominantly adopt an immune-responsive and barrier-supportive phenotype, whereas male ECs show signatures associated with metabolic stress and structural angiogenic remodeling, proteostatic stress adaptation, and extracellular matrix reorganization.

To delineate the specific contribution of YAP/TAZ signaling to these sex-divergent stress responses, we next analyzed endothelial transcriptional and proteomic changes following YAP/TAZ silencing.

### YAP/TAZ silencing reprograms the endothelial stress response

To define the role of YAP/TAZ in endothelial stress adaptation, we performed bulk RNA sequencing and mass spectrometry-based proteomics following siRNA-mediated YAP/TAZ silencing (siYT) in endothelial cells exposed to TNF/Glc, including analysis of the endothelial secretome. TNF/Glc induced a robust inflammatory core response that was largely preserved in both control and siYT endothelial cells, including activation of canonical cytokine and inflammatory pathways (Fig. 5A). However, YAP/TAZ silencing markedly altered the composition of this response. Gene set analysis revealed reduced enrichment of canonical YAP/TAZ target signatures and pathways related to cell cycle, migration, activation, and cytokine production (tmod IDs: LI.M4.1, LI.M122, LI.M24; MSigDB ID: M5897, GL1). Accordingly, expression of YAP/TAZ targets (*CCN1, CCN2, CRIM1, FSTL1*) and endothelial activation markers (*ESM1, EDN1, ANGPT2, CXCLs, IL6*) was attenuated.

**Figure 5.**
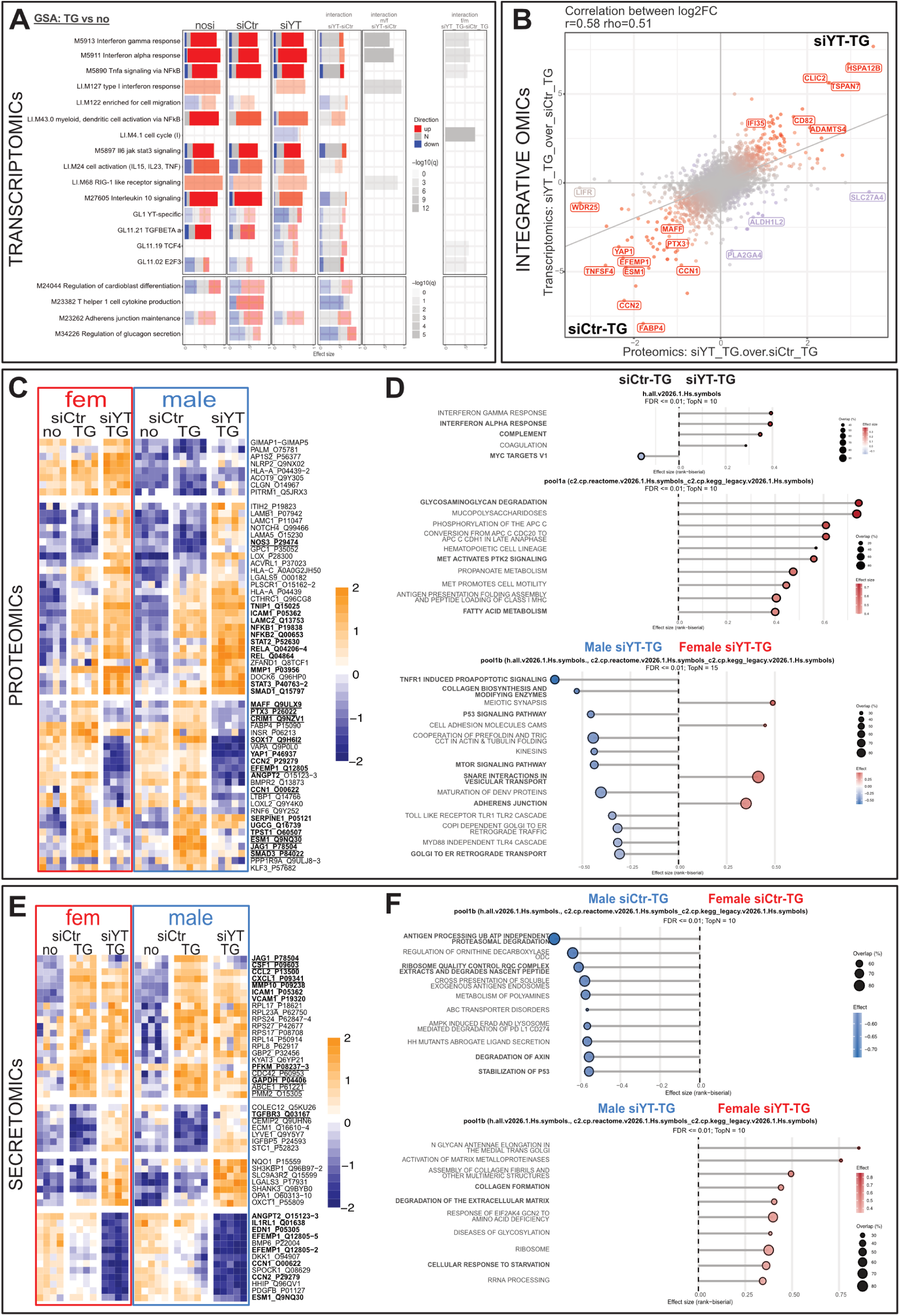
YAP/TAZ signaling controls expression and secretion of HFpEF-associated endothelial biomarkers and sex-specific differential outcome. **(A)** Bulk RNAsequencing results in female and male HUVECs under TNF/Glc (TG) show GSA results for the contrast TG vs no in ECs treated without siRNA (nosi), siCtr or siYAP/TAZ (i.d. no_TG vs no_no; siCtr_TG vs siCtr_no; siYT_TG vs siCtr_no), the interaction (siCtr_TG-siCtr_no)-(siYT_TG-siYT_no), the sex-specific interaction ((fem_siCtr_TG-fem_siCtr_no)-(fem_siYT_TG-fem_siYT_no)-(male_siCtr_TG-male_siCtr_no)-(male_siYT_TG-male_siYT_no)) and the sex-TG-specific interaction ((fem_siYT_TG-fem_siCtr_TG)-(male_siYT_TG-male_siCtr_TG)). Row corresponds to enriched molecular signature database term (tmod, hallmark, kegg, go), bar length indicates effect size (area under curve >0.6), intensity of the color corresponds to the FDR; red and blue fragments indicate fraction of DEGs within the contrast (log2FC>0.5; FDR<0.05) (**A**). GL, gene list. GL1 for YAP/TAZ-specific, GL11.02^81^, GL11.19 TCF4^82^ and GL11.21 TGFBETAa^83^. (**B**) Integrative OMICs results: Discoplot showing the correlation between RNAs and proteins and siYAP/TAZ_TG over siCtr_TG (lower left quadrant= positive correlation in siCtr_TG; upper right quadrant= positive correlation in siYAP/TAZ_TG. (**C-D**) Proteomic analysis of female and male siCtr and siYAP/TAZ endothelial cells (ECs) under TNFGlc (TG) treatment. (**C**) Heatmaps display selected differentially expressed proteins (logFC > 1; adj.P < 0.05). Red: female samples; blue: male samples. (**D**) Dot plot of significantly enriched hallmark, reactome, kegg pathways based on log2 fold changes (siYT-TG vs. siCtr-TG and fem-siYT-TG vs. male siYT-TG; n=5 male, n=4 fem). Enrichment was assessed using a two-sided Wilcoxon rank-sum test with Benjamini–Hochberg correction; pathways with FDR ≤ 0.05 are shown. The x-axis indicates the rank-biserial effect size (positive = higher in WT, negative = higher in dTGR). Point size represents the percentage of quantified proteins overlapping each pathway. (**E-F**) Secretome proteomic analysis of female and male siCtr and siYAP/TAZ ECs under TNFGlc (TG) treatment. Heatmaps display selected differentially secreted proteins (log2FC > 1; adj. P < 0.05). Female: red; male: blue) (**E**). (**F**) Dot plot of significantly enriched hallmark, reactome, kegg pathways based on log2 fold changes (fem-siCtr-TG vs. male-siCtr-TG and fem-siYT-TG vs. male siYT-TG; n=5 male, n=4 fem). Enrichment was assessed using a two-sided Wilcoxon rank-sum test with Benjamini–Hochberg correction; pathways with FDR ≤ 0.05 are shown. The x-axis indicates the rank-biserial effect size (positive = higher in WT, negative = higher in dTGR). Point size represents the percentage of quantified proteins overlapping each pathway.

In contrast, siYT induced a distinct transcriptional program characterized by activation of a type I interferon signature (*STAT2, ISG15, ISG20, OAS3*), BMP/TGFβ signaling, and extracellular matrix and junctional remodeling genes (e.g., *LAMB3, EFNB1*), together with upregulation of the YAP antagonist *VGLL4* (Fig. 5A; Extended Data Fig. 3A,B).

Integration of transcriptomic and proteomic datasets revealed concordant regulation of multiple targets at transcript and protein levels (e.g., PTX3, ESM1), alongside evidence of post-transcriptional regulation for selected genes (Fig. 5B; Extended Data Fig. 3C,D). Proteomic analysis confirmed reduced abundance of endothelial injury-associated and YAP/TAZ-dependent proteins (MAFF, PTX3, EFEMP1, ESM1, CCN1, CRIM1, SOX17, TPST1, JAG1, SMAD3) and increased expression of stress-associated proteins (CLIC2, CD82, IFI35, ADAMTS4) (Fig. 5B,C). Notably, key inflammatory mediators (ICAM1, NFκB1/2, RELA, STAT2/3, MMP1) remained elevated or were further increased under combined siYT and TNF/Glc stress (Fig. 5C).

At the pathway level, 1D enrichment analysis identified enhancement of interferon signaling, glycosaminoglycan degradation, MET/PTK2 signaling, and fatty acid metabolism in siYT ECs under TNFGlc (Fig. 5D, upper panels). Stratification by sex revealed distinct pathway biases: female endothelial cells showed enrichment of adherens junction organization and vesicle transport pathways, whereas male cells were enriched for collagen biosynthesis, TNFR1-mediated apoptosis, p53/mTOR signaling, and Golgi-to-ER transport processes (Fig. 5D, lower panels).

Analysis of the endothelial secretome further demonstrated that YAP/TAZ silencing attenuated the release of multiple injury-associated factors, including CXCL1, CSF1, PFKM, GAPDH, ANGPT2, IL1RL1, EFEMP1, and ESM1 (Fig. 5E). Importantly, secretome profiling uncovered marked sex-specific differences in paracrine signaling. Under TNF/Glc conditions, male endothelial cells preferentially secreted proteins associated with translational machinery, proteasomal degradation, and p53 regulatory pathways. However, YAP/TAZ silencing markedly reduced this anabolic secretory profile in male endothelial cells. In female endothelial cells, siYT instead enhanced secretion of factors linked to extracellular matrix remodeling, collagen formation, matrix degradation, and nutrient stress/starvation pathways (Fig. 5F).

Together, these results show that YAP/TAZ silencing reprograms endothelial stress responses at transcriptional, proteomic, and secretory levels, preserving inflammatory signaling while shifting endothelial output toward a less anabolic, potentially protective state in males, but toward a more remodeling- and stress-associated, potentially disruptive phenotype in females.

To evaluate how these YAP/TAZ-dependent endothelial programs affect the perivascular microenvironment, we next investigated microvascular integrity and pericyte responses under cardiometabolic stress.

### Cardiometabolic stress alters microvascular stability and pericyte responses

A 3D microvascular network-on-chip model was used to examine the effects of cardiometabolic stress on endothelial–pericyte interactions ex vivo^45^. Female or male HUVECs were combined with human pericytes and embedded in fibrin gel within microfluidic chips. After three days of microvascular self-assembly, networks were exposed to sustained metabolic inflammatory stress using TNF/Glc (Fig. 6A).

**Figure 6.**
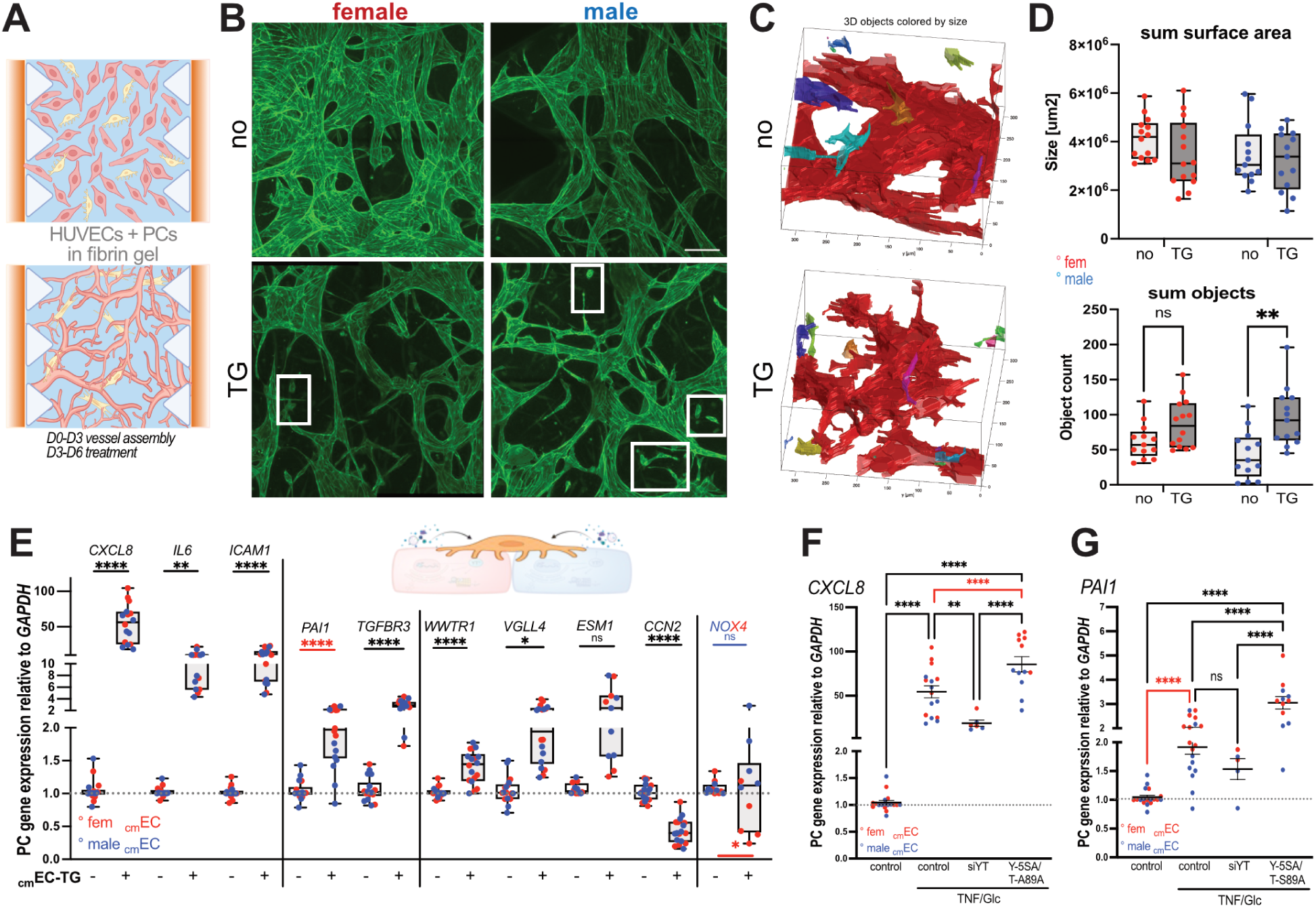
Vascular integrity is compromised in microvessels derived from male ECs under stress: 3D microvascular network and endothelial paracrine effects. (**A**) Schematic representation of the microvascular network-on-chip assay adapted from https://aimbiotech.com using https://biorender.com. After three days of vessel self-assembly, the microvessels were treated for another three days with TNF/Glc (TG). (**B**) Representative immunofluorescence pictures of VE-cadherin (green)-stained microvessels (scale-bar, 100µm) under sustained TG treatment showing increased endothelial sprout rupture (white rectangles) particularly when using male HUVECs in combination with human brain pericytes. (**C**) Representative 3D renderings of the segmented vasculogensis-on-chip assay used for quantifications. Single detected objects are colorcoded. (**D**) Quantifications of microvascularity showing surface area and objects counted (fem-EC-derived, red dots, N=14; male-EC-derived, blue dots, N=13). (**E-G**) Gene expression analysis of pericytes after exposure of conditioned medium derived from female (red) or male (blue) ECs showing increase in the inflammatory and fibrotic response, differential TEAD response and sex-specific differences in the expression of *NOX4* under TG (**E**). (**F,G**) QRT-PCR analysis of pericytes after exposure of conditioned medium from ECs with silenced YAP/TAZ or with transiently transfected YAP-5SA or TAZ-S89A for *CXCL8* (**F**) and *PAI1* (**G**). Data represent N=3-8, analyzed from three independent experiments and displayed as mean ± SEM. One-way analysis of variance (ANOVA) followed by multiple-comparisons test. P values < 0.05 (*), <0.005 (**), <0.0005 (***), <0.0001 (****).

Under TNF/Glc treatment, microvascular networks derived from male ECs exhibited increased microvascular instability and sprout rupture compared to networks derived from female ECs, which maintained greater structural integrity (Fig. 6B; Fig.6C, Extended Data Fig. 4A). Quantitative analysis confirmed increased fragmentation in male networks, reflected by a higher number of disconnected vascular structures (objects) despite unchanged total surface area (sum objects and sum surface area in Fig. 6D). Male networks also contained a higher proportion of small-calibre vessels (1,000–10,000 µm²) arising from larger trunks (Extended Data Fig. 4B).

To assess whether endothelial secreted factors influence the perivascular microenvironment, primary human pericytes were exposed to conditioned medium derived from female or male ECs subjected to cardiometabolic stress. QRT-PCR analysis revealed distinct transcriptional responses depending on the sex of the endothelial donor (Fig.6E). Pericytes exposed to conditioned medium from male ECs increased expression of NADPH oxidase (*NOX4*), whereas *NOX4* expression was reduced following exposure to conditioned medium from female ECs.

In contrast, the pro-fibrotic gene *PAI1* (SERPINE1), was significantly increased in pericytes treated with conditioned medium of female ECs under cardiometabolic stress, while conditioned medium from male ECs had little effect on *PAI1* levels (Fig. 6E). Additional inflammatory and remodeling-associated genes, including *TGFBR3, CXCL8, IL6 and ICAM1* were also altered in pericytes exposed to endothelial conditioned media.

Expression of several YAP/TAZ-associated genes in pericytes was also modulated by endothelial conditioned media, including expression of *WWTR1(TAZ)*, *VGLL4,* and the endothelial YAP/TAZ target *ESM1*, while *CCN2* expression was reduced.

To test whether these effects depend on endothelial YAP/TAZ signalling, endothelial cells were subjected to YAP/TAZ silencing or overactivation prior to conditioned medium transfer. Pericytes exposed to conditioned media from YAP/TAZ-silenced ECs showed reduced expression of *CXCL8* and *PAI1, whereas* conditioned medium from YAP/TAZ-activated endothelial cells increased expression of these genes, particularly *CXCL8* under female EC medium (Fig. 6F,G).

Together, these experiments show that cardiometabolic stress alters microvascular stability and pericyte responses in a sex-dependent manner and that these effects are influenced by endothelial YAP/TAZ signaling.

Across human plasma profiling, mouse genetics, endothelial signaling analyses, and microvascular modelling, these findings identify endothelial YAP/TAZ signaling as a central regulator linking cardiometabolic stress to sex-divergent endothelial and microvascular remodeling in HFpEF.

## Discussion

HFpEF remains a therapeutic challenge, in part due to its pathophysiological heterogeneity and lack of mechanistically informed biomarkers or treatment targets. In this study, we show that endothelial YAP/TAZ signaling integrates cardiometabolic stress responses and contributes to sex-divergent endothelial and microvascular remodeling in HFpEF. By combining human plasma proteomics, inducible endothelial mouse genetics, sex-stratified endothelial multi-omics, and microvascular organ-on-chip modeling, our findings link circulating YAP1 signatures with endothelial stress-induced uncoupling of YAP/TAZ-TEAD activity, which reshapes the endothelial secretome and propagates distinct perivascular remodeling programs in males and females.

Our analysis of UK Biobank data suggests that YAP1 is a circulating biomarker associated with HF and adverse outcomes. Importantly, YAP1 was not uniformly increased in obesity or diabetes alone, suggesting that its elevation reflects manifest endothelial dysfunction rather than cardiometabolic risk *per se*. Notably, YAP1 is primarily known as an intracellular transcriptional co-activator and is not typically considered a secreted protein. Its detection at measurable levels in human plasma therefore suggests that cardiometabolic/hypertensive stress may lead to extracellular release of YAP1, potentially through cell injury, vesicle-mediated export, or other stress-associated mechanisms. Parallel increases in endothelial activation markers, including ESM1, ANGPT2, EDN1, VCAM1, and CDH5, support an endothelial origin of circulating YAP1 and implicate microvascular activation as a central component of HFpEF pathobiology. Coronary microvascular dysfunction and endothelial inflammation are hallmarks of HFpEF and precede overt myocardial remodeling^46–48^. ANGPT2-mediated vascular destabilization, EDN1-driven vasoconstriction, and inflammatory cytokines such as IL-6 and PTX3 have all been linked to HFpEF severity and adverse outcomes^49–53^. Our findings position YAP1 within this endothelial injury signature and suggest that circulating YAP1 may serve as a biomarker of microvascular stress and prognostic risk, particularly in male HFpEF patients. We also detected YAP1 in male EC culture medium, with levels increasing following TNF/Glc treatment. Whether YAP1 is actively secreted or released during endothelial stress remains to be determined.

In the hypertensive cardiomyopathy and nephropathy mouse model, endothelial-specific YAP/TAZ deletion preserved systolic function, prevented adverse ventricular remodeling, and improved global longitudinal strain compared with controls and TAZ gain-of-function mice. These findings are consistent with restored endothelial–myocardial coupling, a concept central to HFpEF pathogenesis^46,54^. Endothelial dysfunction reduces nitric oxide bioavailability, promotes inflammation, and impairs cardiomyocyte relaxation and contractility^21–24^. By attenuating the maladaptive endothelial secretome, YAP/TAZ deletion likely improves microvascular perfusion and paracrine support of cardiomyocyte function. Notably, sex-specific outcomes were observed: fibrosis was reduced predominantly in female YAP/TAZ-deficient mice, whereas survival benefit was more pronounced in males. This divergence mirrors the in vitro findings. Female endothelial reprogramming favoured matrix modulation and TGFβ buffering, consistent with reduced fibrotic remodeling. Male endothelial reprogramming enhanced stress resilience and reduced profibrotic and proinflammatory paracrine cues, potentially mitigating lethal decompensation under combined cardiac and renal stress. Collectively, these data position endothelial YAP/TAZ signaling as a critical regulator of coronary microvascular integrity and secretome-driven myocardial remodeling under cardiorenal stress. This aligns with previous reports implicating YAP1 in endothelial inflammation and vascular stiffening^16,55^, but our work extends this by demonstrating causality in the HFpEF context and highlighting sex-specific susceptibility, particularly the vulnerability of male mice to YAP/TAZ activation.

Mechanistically, cardiometabolic stress (glycolytic constraint or TNF/Glc) suppressed canonical YAP-TEAD transcriptional activity while activating NFκB, and BMP/TGFβ pathways, suggesting a qualitative rewiring of YAP/TAZ signaling. YAP/TAZ are well-established mechanotransducers that couple cytoskeletal tension and metabolic cues to transcriptional programs governing proliferation, survival, and angiogenesis^2–4^. However, their role under combined inflammatory and metabolic stress appears more complex. Under glycolytic constraint, male ECs may engage alternative TEAD cofactors (e.g., *VGLL3*, *TEAD3*) and KLF4-dependent programs, suggesting a shift from canonical YAP-TEAD signaling toward a non-canonical transcriptional axis associated with endothelial phenotypic plasticity and early EndMT-like states. In contrast, female ECs showed stronger suppression of TEAD reporter activity without comparable compensatory cofactor induction. Under TNF/Glc, YAP was partially uncoupled from nuclear TEAD activity, with induction of the competitive inhibitor VGLL4^40^ and increased extracellular YAP1 release, particularly in males. These findings indicate that cardiometabolic inflammation qualitatively reshapes the transcriptional output of the endothelial YAP/TAZ pathway in a sex-dependent manner.

Sex differences in endothelial biology, immune responses, and HFpEF prevalence are well established (reviewed in^56^). Our data extend these observations by demonstrating that YAP/TAZ signaling operates within a sex-specific regulatory network that determines endothelial adaptation to metabolic stress. Bulk transcriptomic and proteomic profiling under TNF/Glc identified a shared inflammatory core in female and male ECs, marked by interferon signaling, NFκB activation, and cytokine pathway enrichment. This profile is consistent with the inflammatory HFpEF paradigm^46,47,54,57^. However, downstream remodeling programs diverged substantially.

Female ECs adopted an immune- and barrier-responsive phenotype characterized by interferon signaling and regulated extracellular matrix turnover, consistent with adaptive remodeling that preserves vascular integrity. In contrast, male ECs displayed a metabolically strained and structurally unstable phenotype, marked by altered lipid handling, proteostatic stress, and features of endothelial plasticity. Functionally, male-derived microvascular networks exhibited greater thinning and sprout rupture under cardiometabolic stress, consistent with unstable angiogenesis and microvascular rarefaction^24,57–59^. Notably, YAP/TAZ silencing attenuated endothelial injury-associated secretome factors (including IL6, EDN1, ANGPT2, PTX3, CSF1, CXCL chemokines, and ESM1) without abolishing inflammatory signaling, but further amplifying interferon type I programs, indicating that the core inflammatory response to cardiometabolic stress is largely independent of YAP/TAZ signaling. Instead, YAP/TAZ appears to modulate the organization and downstream integration of this response. This modulation was sex-specific: female YAP/TAZ-deficient ECs disrupted the stress-adaptive program and shifted the endothelial state toward maladaptive remodeling. Hallmarks of this shift were collagen-associated pathways linked to extracellular matrix remodeling, and nutrient stress/starvation pathways, suggesting loss of endothelial identity and barrier integrity. In contrast, male YAP/TAZ-deficient ECs shifted from a secretory proteostatic towards a more quiescent protective state. Together, these findings position endothelial YAP/TAZ as a regulator of the inflammatory secretome’s composition and of sex-dependent microvascular remodeling trajectories under cardiometabolic stress. This intrinsic sexual dimorphism in endothelial responses is further supported by our 3D microvascular network-on-chip assays, where microvessels derived from male ECs showed greater vascular instability and fragmentation under metabolic inflammatory stress. Moreover, secreted proteins, including YAP1, from stressed ECs exerted sex-specific paracrine effects on pericytes, suggesting an additional level of vascular destabilization mediated by EC-secreted factors. Together, these findings indicate that endothelial YAP/TAZ functions both as an intracellular integrator of biomechanical and metabolic stress and as a regulator of paracrine signaling that shapes microvascular stability. The functional relevance of this endothelial reprogramming was supported *in vivo* using a hypertensive cardiomyopathy and nephropathy disease model, where endothelial YAP/TAZ deletion restored ejection fraction, reduced left ventricular systolic dimensions, and enhanced systolic wall thickening. These improvements were accompanied by sex-specific outcomes: fibrosis rescue was most pronounced in females, whereas improved survival was observed predominantly in males. Together, these findings indicate that YAP/TAZ deletion re-establishes endothelial–myocardial coupling by reducing microvascular destabilization and restoring microvascular support of cardiomyocyte contractile function, while engaging distinct protective pathways in female versus male endothelium.

Overall, these data position endothelial YAP/TAZ as a key molecular switch that converts cardiometabolic stress into sex-specific maladaptive endothelial states, whereas its inhibition promotes microvascular stabilization, improved myocardial performance, and divergent but complementary protective outcomes in females and males.

From a translational standpoint, our findings carry several important implications. First, circulating YAP1, particularly in combination with ESM1, may serve as a biomarker of endothelial stress and prognostic risk in HFpEF. Its detectability in human plasma and predictive association with mortality underscore its potential utility as a stratification tool, especially in identifying endothelial-dominant HFpEF phenotypes. Second, the sex-divergent rewiring of YAP/TAZ signaling suggest that endothelial responses to cardiometabolic stress differ between males and females, supporting the development of sex-stratified therapeutic strategies. Third, the switch from canonical TEAD1/4 to TEAD3 signaling and alternative cofactors VGLL3/4 may offer opportunities to modulate endothelial plasticity, perhaps allowing selective rebalancing of maladaptive responses without global inhibition of YAP/TAZ.

### Limitations and Future Directions

While our study integrates human data, mouse models, and mechanistic cell biology, several limitations should be noted. First, the precise tissue source of circulating YAP1 in humans remains to be definitively established, although our data strongly point to endothelial origins. Second, hormonal regulation of the observed sex differences, particularly the role of estrogen, was not directly interrogated here and will require future exploration. Third, while we focused on HFpEF, some of the pathways identified may be relevant to other comorbidities (e.g., diabetes, obesity) and merit broader investigation.

## Conclusion

This study positions endothelial YAP/TAZ signaling as a mechanistic and biomarker axis in HFpEF pathogenesis. We reveal that under cardiometabolic stress, ECs undergo sex-specific transcriptional rewiring, leading to maladaptation in males and relative resilience in females. Circulating YAP1 emerges as a promising marker of this process and may help stratify HFpEF patients for future targeted interventions. Our work opens new avenues for sex-aware, endothelium-focused diagnostics and therapies in heart failure.

## Material and Methods

### Human Plasma Proteomics Analysis

#### UK Biobank disease and control group selection

Several clinical subgroups were defined within the UK Biobank cohort based on diagnostic codes and phenotypic data. These included individuals classified as heart failure with preserved ejection fraction (HFpEF, n = 33,480), diagnosed hypertension (aHTN, n = 40,402; based on ICD-10 codes), and heart failure with reduced ejection fraction (HFrEF, n = 486). Control groups were also identified: a large healthy group with no history of heart failure diagnosis or symptoms (non-HF, n = 241,611), as well as diabetic controls (n = 9,048) and obese controls (n = 36,455). Full inclusion and exclusion criteria for these groups are described in a recent publication^25^. Proteomics analyses were restricted to the subset of participants with available YAP1 expression data, which represented approximately 10% of the full cohort. This resulted in the following group sizes: HFpEF (n = 2,712), Non-HF(n = 20,600.), aHTN (n = 3,911), diabetic (n = 809), obese (n = 2,884), and HFrEF (n = 43).

#### Confounding analysis

To compare protein expression levels and assess confounding effects of covariates, including medication intake, comorbidities, and lifestyle factors, the *metadeconfoundR* package (v.0.3.0) was used. This method comprised two main steps. First, the naïve associations between omics features and both the disease group of interest and covariates are computed. Second, confounding is assessed using post hoc nested linear model comparison via likelihood ratio tests (LRT). The association is considered deconfounded if the disease status remains significantly associated with the omics feature after adjusting for the covariate (LRT < 0.05). This means that disease status adds explanatory power beyond the tested covariate in the specified nested model. Conversely, if the covariate contributes more to the association than the disease status itself, the association is labelled confounded (Forslund et. al., 2021). The analysis was conducted independently for each disease vs. control comparison (each column in Figure 1A), covering 8 proteins and 131 covariates.

#### Kaplan-Meier survival curves

Kaplan–Meier survival curves were generated to assess overall survival across subgroups defined by varying levels of YAP1 expression (Figure 1B). Survival analysis was performed using the *lifelines* Python library. The event of interest was all-cause mortality, and the time variable was defined as the number of years between the recruitment date and either the recorded date of death or the censoring date. Participants who had no recorded death as of November 16, 2023, were considered right-censored at that date. The analysis was stratified by clinical and demographic subgroups, including sex, heart failure status (non-HF or HFpEF), and the presence or absence of hypertension. Within each stratified group, participants were further categorised into four groups based on YAP1 expression levels: below the 25^th^ percentile, between the 25^th^ and 75^th^ percentiles, between the 75^th^ and 95^th^ percentiles, and above the 95^th^ percentile. Kaplan-Meier survival curves were estimated separately for each YAP1 expression group within each stratified subgroup.The statistical significance of survival differences between groups was evaluated using the log-rank test, applied to all pairwise comparisons within each stratification. Confidence intervals at the 95% level were included to indicate uncertainty in the survival estimation.

### Genetic mouse models and treatments

For loss and gain of function experiments the following mouse strains in C57BL/6J background were used: Yap fl/fl and Taz fl/fl^60^, Taz GOF (3xFLAG-TAZS89A-IRES-nEGFP^3,4^, and Pdgfb-iCreERT2^61^.

All mouse experiments complied with the German Animal Protection Act and were approved by the local Berlin authority, the Landesamt für Gesundheit und Soziales (LaGeSo, TVV281-19). Mice were maintained at the Max Delbrück Center for Molecular Medicine under standard husbandry conditions (22 ± 2 °C, 55 ± 10% humidity, 12:12 h light–dark cycle). Tamoxifen (Sigma) was injected intraperitoneally (IP; 100µg/g body weight) in 7-9 weeks-old mice every other day for a total of three times. Animal experiments in mice were performed using an established cardiorenal hypertension protocol (Marko et al., 2020; Tsukamoto et al., 2013) of combined uni-nephrectomization with Angiotensin II (AngII)-administration via osmotic minipumps (1.44 mg/kg/day by subcutaneous osmotic minipumps, Alzet, size 2004). The investigators were blinded to allocation during experiments.

### Echocardiography and cardiac fibrosis assessment

For echocardiography on anesthetized 13-15-week-old mice, the Vevo 770 system (Visual Sonics, Inc.) with a 45 MHz transducer mounted on an integrated rail system was used. Standard imaging planes and functional calculations were obtained according to the American Society of Echocardiography guidelines. The LV parasternal long axis 4-chamber view was used to derive fractional shortening (%FS), ejection fraction (%EF), and ventricular dimensions and volumes. Myocardial strain indicative of the diastolic function was quantified by two-dimensional speckle-tracking echocardiography using high-frame-rate images acquired in parasternal long- and short-axis views. Strain analysis was performed offline by an investigator blinded to group allocation.

Myocardial samples were fixed in 10% paraformaldehyde and embedded in paraffin. Sections of 5 μm thickness were stained with Masson-Goldners trichrome staining (Carl Roth, #3459.1) according to the manufacturer’s instructions for visualization of extracellular matrix deposition. Stained slides were dehydrated, cleared, and mounted with a Eukitt Quick-Hardening mounting medium (Merck KG, #03989). Collagen fibers appeared blue/green, whereas cytoplasm and muscle elements stained red, providing reliable structural contrast for the assessment of interstitial fibrosis. Internal staining controls were included in each run to ensure reproducibility. Presence or absence of myocardial fibrosis was assessed in one section per animal by a blinded physician scientist experienced in the interpretation of myocardial histology. Depending on the localization of the lesions, myocardial fibrosis was categorized as perivascular, subepicardial or diffuse.

### Cell culture and stress modelling

HUVECs from pooled donors (PromoCell; pooled from single-female donors: C-12200, 479Z024, 485Z034, 488Z021, 433Z035.1 ; male-pool: C-12203, 479Z016) and human brain pericytes (ScienCell, Reference: #1200, 27194, single donor) were cultured on 0.2% gelatine precoated dishes/flasks in EGM2-Bulletkit without antibiotics (Lonza, #CC-3156) or in pericyte medium (PM, ScienCell, #1201), respectively.

For knockdown experiments, HUVECs were transfected with SMARTpool: siGENOME siRNAs purchased from Horizon (Yap #M-012200-00-0005, Taz #M-016083-00-0005, and non-targeting siRNA Pool 1 #D001206-13-05). Briefly, subconfluent (70-80%) HUVECs were transfected with 25 nM siRNA using Dharmafect 1 transfection reagent following the protocol from the manufacturer; transfection media was removed after 24 hours, cells were transferred to 80rpm orbital shaking and experiments were routinely performed on the third day after siRNA transfection.

To activate YAP, TAZ and VGLL4 signalling in ECs, female and male HUVEC pools were transfected with YAP-5SA (Addgene #33093); YAP-5SA,S94A (Addgene #33103), TAZ-S89A (Addgene #32840) or VGLL4 (SinoBiological #HG18349-UT) plasmids using TransIT 2020 (Mirus), pcDNA3.1 was used as a control. Transfections were carried out by incubating sub-confluent HUVECs (70-80%) with starvation media (DMEM containing 2% FBS) for 4 hours. After 3 hours, transfection media was removed and cells were cultured in complete EGM2 media (Lonza) while orbital shaking (80rpm). All experiments were performed 48 hours post transfection.

For the induction of metabolic stress responses in HUVECs, transfected and nontransfected HUVEC pools were treated for 17 hours with 2-Desoxy-D-glucose (1mM or 10mM 2DG, Sigma D6134) or TNFa and Glucose (5 or 10ng/ml TNF; Gibco, #PHC3015; 0.5% D-(+)-Glc; Sigma, #G8769), respectively.

For the generation of OMICs, ELISA and EC-pericyte co-culture data, 55 hours post siRNA transfection or 31 hours post plasmid transfection female and male HUVECs were cultured in phenol-free EGM-Bulletkit without FBS, with antibiotics (LONZA) and the respective treatment on an orbital shaker (INFORS HT Celltron, #79330) at 80rpm. After 17 hours, cells were frozen at -80 degree for RNAseq experiments or collected in cold PBS, centrifuged for 5min at 800g and frozen in 50 ul PBS for mass spectrometry. The supernatant was collected and centrifuged for 5 min at 300g and either supplied to pericytes or centrifuged a second time for 5 min at 17.800g and frozen for mass spectrometry or used for human YAP1 ELISA (MyBioSource, #MBS7238166) experiment according to the manufacturer’s instructions.

### Dual luciferase reporter assay

Renilla-luciferase reporter assays for TEF-1 (TEAD signaling^62^), BRE (Bmp signaling^63^), CAGA (TGFß signaling^64^, NFkB^65^ (pGL2-basic-HindIII-IgK(3x)cona-pBluescriptSKII-BamH1) and FOPflash^66^-Luciferase promoter activity were performed as follows: 48 hours after gene knockdown by siRNA or 24 hours after plasmid overexpression HUVECs were cotransfected with 600 ng of Luciferase reporter gene construct and 300 ng of pRL-CMV (Promega) using Lipofectamine2000 and incubated for 3 hours. Cell extracts were prepared 24 hours post Luciferase reporter transfection, and luciferase activity was measured using a dual luciferase system (BertholdTech CENTRO, Driver Version: 1.21, (1.0.21.0), S/N: 50-6902, Embedded Version: 2.08) as described^67^. Experiments were carried out in duplicates and results were normalized to the corresponding FOPflash/Renilla measurement. For quantitative analyses, a minimum of three biological replicates were analysed.

### RNA extraction and quantitative real time-polymerase chain reaction

RNA was extracted using the Nucleo Spin RNA Mini Kit (Macherey-Nagel, #740933.250) according to the manufacturer’s instructions. For HUVECs transfected with siRNAs or plasmids, 90 ng of RNA were reverse transcribed using RevertAid First Strand cDNA Synthesis Kit (ThermoFisher Scientific, #K1622). qRT-PCR was performed using TaqMan reagents and probes (Applied Biosystems) (listed in Suppl.Tab.S1). qRT-PCR reactions were run on Quant Studio 6 Flex (Applied Biosystems) and gene expression was calculated with the comparative 2deltaCT method and normalised to human GAPDH.

### Immunofluorescence staining and antibodies

For immunofluorescence in HUVECs, cells were grown in #1.5 coverslips coated with poly-lysine (Merck, #P4822) and gelatin 0.2% (Merck, #G1393). At the end of the experiment cells were fixed in 4% PFA (Merck, #158127) for 10 min, permeabilised in 0.3% Triton-X100 in blocking buffer for 5min and blocked in 1% BSA 20mM Glycine in PBS for 30 min. Primary and secondary antibodies were incubated for 2 and 1 hours in blocking buffer, respectively. Nuclei labeling was performed by incubating cells with DAPI for 5 min (Life technologies, #D1306).

Aimbiotech devices were fixed with 4% PFA for 20 min, blocking and permeabilization were performed using blocking buffer containing 3% BSA (Serva, #11930.03), 0.05%Triton X-100, 0.01% sodium deoxycholate (Merck, #D6750-25G), and 0.02% sodiumazide (Merck, #S2002-25G). Primary and secondary antibodies were incubated for 72 and 24 hours in blocking buffer at 4°C, respectively. Nuclei labeling was performed by incubating devices with DAPI for 5 min followed by postfixation in 2% PFA. Devices were imaged on the Leica MICA microscope for overview images (10x objective) and on the Zeiss LSM 980 for high-resolution confocal imaging (40x objective). Z-stacks were acquired in confocal mode to capture the full 3D vascular network, using a z-step size of 1 µm distance. Maximum-intensity projections of z-stacks were generated with ImageJ for visualization.

A list of the primary and secondary antibodies used can be found in Suppl.Tab.S2.

### 3D Microvascular network-on-a-chip assay, image processing and segmentation

The microvascular network-on a-chip assay was performed in a commercially available 3D cell culture microfluidic chip (AIMBiotech, idenTx #DAX-1) using the 2-step seeding method as described in^45^. Briefly, after trypsinization HUVECs and pericytes were resuspended in thrombin (Merck, #T4393-100UN), stock reconstituted in water and diluted in EBM-2 (Lonza, #CC-3156) to a final concentration of 2 U/mL at a cell concentration of 20 × 10⁶ cells/mL. For step one, coating ECs along the micropillars, HUVECs-thrombin solution was mixed 1:1 with fibrinogen (5 mg/mL; Sigma, #F3879), flashed and aspirated from the middle gel chamber of the AIM Biotech chip. For the second seeding step, HUVECs were mixed with human brain pericytes at a 1:4 ratio and combined with fibrinogen before seeding the mixture into the same middle gel chamber. After polymerization of the fibrin–thrombin–cell suspension, side channels were gelatinized (0.2% gelatin; Merck, #G1393). Side channels were then seeded with 10 µL of brain pericytes at 0.4 × 10⁶ cells/mL. Chips were incubated at 37°C and 5% CO₂ in EGM-2 supplemented with VEGF (Pepro Tech, #100-20-250UG) to a final concentration of 50 ng/mL Medium was changed every second day. For inflammatory/glucose stress treatment, on day three and five of culture, EGM2 with VEGF with or without TNF-α (3,3 ng/mL; Gibco, #PHC3015) and glucose (0.5% D-(+)-Glc; Sigma, #G8769) were added. On day six vascular networks were fixed, followed by immunostaining and imaging on the Leica MICA microscope. Raw microscopy data were processed using Fiji (ImageJ, National Institutes of Health, Bethesda, MD) to enable analysis of similarly sized regions located between the AimBiotech pillars. The cropped regions were then segmented, and quantification was performed using a custom-written high-throughput MATLAB (version 2025b) pipeline. Briefly, to account for image-to-image background heterogeneity, each z-slice was flat-field^68^. The corrected images were subsequently subjected to 2D adaptive noise-removal filtering using a low-pass Wiener filter. Signal segmentation was then performed using the Difference of Gaussian (DoG) method, as previously described^69^. Gaps in the segmentations were closed through a sequence of mask dilations and erosions, applied first in 2D and subsequently in 3D. The resulting signal mask was used for downstream data visualization and analysis, including 3D rendering (Fig.6C), maximum-intensity projections (Extended Data Fig. 3A, top), quantification of the surface area of segmented objects (Extended Data Fig. 3B), and analysis of object-to-object connectivity (Extended Data Fig. 3A, bottom).

The code generated to analyze the data in this publication will be made available upon publication. The code can be shared with the reviewers if requested. Code will be deposited on github.

### Mass spectrometry data acquisition and analysis

Cells and conditioned media were lysed in sodium deoxycholate (SDC) buffer, heat-denatured (95 °C, 10 min), reduced with dithiothreitol (10 mM), and alkylated with chloroacetamide (40 mM). Proteins were digested overnight at 37 °C using Lys-C and trypsin (1:50 enzyme-to-protein ratio). Digestion was stopped by acidification (1% formic acid), precipitating SDC. Peptides were desalted (AssayMAP Bravo, Agilent), dried, and reconstituted in 3% acetonitrile/0.1% formic acid.

Peptides (1 µg) were separated by reversed-phase LC (Vanquish Neo, Thermo Fisher) on a 20 cm in-house packed C18 column using a 56-min gradient at 0.25 µL/min, and analyzed on an Exploris 480 mass spectrometer in data-independent acquisition (DIA) mode with 12 m/z staggered windows. MS1 scans were acquired at 120,000 resolution (m/z 350–1,650) and MS2 scans at 30,000 resolution using stepped collision energies.

Raw data were processed with DIA-NN (v1.9)^70,71^ using a UniProt human database (including isoforms and contaminants) and a precursor-level FDR of 0,01. Match-between-runs was enabled, allowing one missed cleavage. Protein inference was performed at gene level, and label-free quantification (LFQ) values were derived from maximum precursor intensities per run, protein group and gene.

Data analysis was conducted in R (v4.4.1). Log2-transformed LFQ intensities were filtered (≥3 valid values per group; optional ≥2 peptides per protein), and missing values were imputed using a down-shifted Gaussian distribution. Differential abundance was assessed using limma^72^ with Benjamini–Hochberg correction (adjusted P < 0.05).

Pathway-level changes were evaluated using one-dimensional annotation enrichment^73^. Protein groups were collapsed to gene-level values by retaining the maximum absolute log2 fold change per gene (GCT format). Gene set (GMT format) enrichment (e.g., MsigDB collections) was tested using two-sided Wilcoxon rank-sum tests (minimum 5 genes per set), with effect sizes reported as rank-biserial correlations. Effect sizes were calculated as rank-biserial correlations, derived from the Mann–Whitney U statistic^74^. Specifically, ranks were computed across the combined foreground and background distributions, the Wilcoxon rank-sum statistic (W) was calculated, converted to U, and scaled to the interval [-1, 1]. Positive values indicate higher abundance in the numerator condition, whereas negative values indicate higher abundance in the denominator condition.

P values were adjusted using Benjamini–Hochberg correction, and FDR-significant gene sets were considered enriched.

Proteomic and secretomic 1D annotation enrichment output files can be found in Suppl.Tab.S3A, B.

### RNA sequencing and gene set enrichment analysis

RNA was extracted using the Nucleo Spin RNA Mini Kit (Macherey-Nagel, #740933.250) according to the manufacturer’s instructions. RNA quality has been evaluated by Qubit/TapeStation QC (#QC103X/ #QC101X). RNA libraries were prepared from total RNA using NEBNext Poly(A)mRNA Magnetic Isolation Module (Human/Mouse/Rat) (NEB, #E7490) and NEBNext® Ultra II Directional RNA Library Prep Kit for Illumina® Version 4.0_4/21 (NEB, #E7760L). RNAseq was done on an Illumina NovaSeq X Plus platform in a 100+10+10+100 nt paired-end mode (Illumina). Sequences were mapped to human genome, GRCh38 p7 with GENCODE annotation v. 25 using the STAR aligner v. 2.7.11a^75^ and read counts were summarized with featureCounts v. 2.0.3^76^. Differential gene expression analysis was performed with the R package DESeq2, v. 1.38^77^, and gene set enrichments were obtained through the R package tmod 0.50.13^78^ on built-in tmod transcriptional modules and using gene sets from the MSigDB^79^ as provided by the R package msigdbr, v. 7.5.1^80^. P-values were adjusted to False Discovery Rate (FDR) using the Benjamini-Hochberg procedure.

### Statistics and Reproducibility

For quantitative analyses, a minimum of three biological replicates were analysed. Images from immunofluorescence studies are representative of the respective phenotype observed in samples from at least three independent experiments. Statistical analyses were performed by one-way ANOVA with Tukey’s multiple comparisons test, for luciferase reporter analysis by two-sided Wilcoxon t-test (paired, nonparametric), expcept for luciferase reporter analysis under treatment and overexpression of plasmids, where an ordinary one-way ANOVA with Tukey’s multiple comparisons test was used; and for Yap1 endothelial release data by two-sided Mann-Whitney t-test (unpaired, non-parametric). Microvascular network-on-chip size distributions and object count were assessed for normality using the Shapiro–Wilk test, group differences were evaluated using 2-way ANOVA, followed by Tukey’s multiple comparisons test. For all bar graphs, data are represented as mean ± s.e.m. A value of P < 0.05 was considered significant. Calculations were performed using the Prism v.10.6.1 software (GraphPad Software Inc.).

## Data availability

RNAseq data sets have been deposited in the National Center for Biotechnology Information Gene Expression Omnibus with the accession number GSE320481. Proteomics data sets will be deposited in the PRIDE Archive (Proteomics IDEntifications Database) Source data are provided with this paper. All other data supporting the findings of this study are available from the corresponding author upon reasonable request.

## Use of large language models (LLMs)

Large language models (LLMs), including ChatGPT (OpenAI), were used to assist with non-scientific aspects of manuscript preparation, including rephrasing existing text for clarity and readability.

## Affiliations

Integrative Vascular Biology Laboratory (A.K.B., K.K., T.N., E.B.K, K.M., I.H., H.G.), Experimental and Clinical Research Center (L.S., I.K., D.N.M.), Angiogenesis & Metabolism Laboratory (M.P.), Animal Phenotyping (M.T., A.H.), Translational Bioinformatics (J.W., D.B.) and Proteomics Platform (O.P., P.M.), Max-Delbrück Center for Molecular Medicine in the Helmholtz Association (MDC), Berlin, Germany. DZHK (German Centre for Cardiovascular Research), partner side Berlin, Germany (A.K.B., K.K., T.N., E.B.K., K.M., I.H., U.L., P.M., M.K., D.N.M., H.G.). Medical Department IV-Nephrology and Hypertension (L.S.), UKSH, Kiel, Germany. Institute of Computer-assisted Cardiovascular Medicine (J.V., M.K.), Deutsches Herzzentrum der Charité, Berlin, Germany. Deutsches Herzzentrum der Charité, Department of Cardiology, Angiology and Intensive Care Medicine (U.L.), Campus Benjamin Franklin, Berlin, Germany. Department of Congenital Heart Disease – Pediatric Cardiology (M.K.), Deutsches Herzzentrum der Charité, Berlin, Germany. Friede Springer Cardiovascular Prevention Center (U.L.) at Charité - Berlin, Germany. Helmholtz Institute for Translational AngioCardiosciences (HI-TAC) (H.G.), Max Delbrück Center for Molecular Medicine at Heidelberg University, Heidelberg, Germany. Charite-Universitätsmedizin, Berlin, Germany (J.V., P.M., M.K., D.N.M., H.G.). Max-Delbrück Center for Molecular Medicine in the Helmholtz Association (MDC), Berlin, Germany (D.N.M.). Charité-Universitätsmedizin Berlin, corporate member of Freie Universität Berlin and Humboldt-Universität zu Berlin, Berlin, Germany (D.N.M.). BIH/MDC Genomics Technology Platform, Berlin, Germany (T.B.). Berlin Institute of Health (BIH), Germany (J.W., T.B., D.B., M.P., U.L., H.G.).

## Author Contribution

A.K.B.: Conceptualization, Resources, Data curation, Formal analysis, Funding acquisition, Validation, Investigation, Visualization, Methodology, Writing—original draft, Project administration, Writing—review and editing. L.S.: Data curation, Formal analysis, Funding acquisition, Validation, Investigation, Visualization, Methodology, Writing—review and editing. J.V.: Data curation, Formal analysis, Validation, Investigation, Visualization, Methodology, Writing—review and editing. K.K.: Data curation, Investigation, Methodology, Writing—review and editing. T.N.: Data curation, Formal analysis, Validation, Visualization, Methodology, Writing—review and editing. E.B.K.: Data curation, Methodology. O.P.: Formal analysis, Validation, Investigation, Visualization, Methodology, Writing—review and editing. J.W.: Formal analysis, Validation, Investigation, Visualization, Methodology, Writing—review and editing. K.M.: Data curation, Methodology. I.H.: Data curation, Methodology. I.K.: Data curation, Methodology. M.T.: Data curation, Methodology. A.H. Data curation, Methodology. T.B.: Data curation, Funding acquisition, Methodology. D.B.: Data curation, Funding acquisition. M.P.: Resources, Writing—review and editing. U.L.: Writing—review and editing. P.M.: Funding acquisition, Methodology, Writing—review and editing. M.K.: Conceptualization, Data curation, Formal analysis, Validation, Investigation, Visualization, Methodology, Writing—review and editing. D.N.M.: Conceptualization, Funding acquisition, Writing—review and editing. H.G.: Conceptualization, Resources, Funding acquisition, Writing—original draft, Project administration, Writing—review and editing.

## Acknowledgements

We thank all members of the integrated Vascular Biology laboratory and J. Heinecke for helpful discussions and comments. We thank A. Behrens for kindly providing the *Taz fl/fl* mice. We also thank M-B. Köhler for technical assistance in tissue slicing and staining, C. Janetzki for the library preparations for RNA sequencing and M. Haji for sample preparation for mass spectrometry. We thank W. Birchmeier, D. Besser, E. Sahai, N. Tapon and R. Schmidt-Ullrich for the generous gift of the luciferase plasmids; and N. Reuther and J. Schwarzkopf for help in setting up the microvascular network-on-chip assay in our lab.

## Extended Data figure legends

**Extended Data Figure 1. Circulating ESM1 is decreased in HFpEF with aHTN and correlates with a better survival rate than in HFpEF-non aHTN patients**. (**A**) Heatmap showing the association of YAP1-TEAD, endothelial and HF-associated proteins in plasma of UKB patients (HFpEF aHTN vs HFpEF non-aHTN) using Olink3000 and was tested for potential confounding variables: circles (o) denote confounding features and asterisks (*) indicate deconfounded features. Statistical significance is indicated as follows: false discovery rate (FDR): <0.1 (*), <0.01 (**), <0.001 (***), empty cells represent FDR ≥0.1. (**B**) Kaplan-Meier survival curves stratified by YAP1 expression levels and clinical subgroups in men and women. Each curve represents one of four YAP1 expression ranges: below the 25_th_ percentile (blue), within the interquartile range (25_th_ -75_th_ percentile, orange), between the 75_th_ and 95_th_ percentile, and above the 95_th_ percentile. Columns correspond to clinical groups: HFpEF non-aHTN and HFpEF with aHTN. Rows represent stratification by sex (female on top, male on bottom). Shaded regions indicate 95% confidence intervals.

**Extended Data Figure 2. Endothelial YAP/TAZ loss protects from cardiac fibrosis particularly in females under cardiorenal hypertensive stress.** (**A**) Quantification of cardiac fibrosis subtypes (subepicardial, light blue bars; perivascular, red bars; diffuse, dark blue bars). (**B**) Representative pictures of fibrosis staining (red, cytoplasm/muscle; blue/green, collagen). Data displayed as mean ± SEM (fem, red; male, blue). One-way analysis of variance (ANOVA) followed by multiple-comparisons test. P values < 0.05 (*), <0.005 (**), <0.0005 (***), <0.0001 (****).

**Extended Data Figure 3. YAP/TAZ signaling controls expression of HFpEF-associated endothelial biomarkers and sex-specific differential outcome**. (**A**) Evidence plots of GO terms representing differential gene expression regulation under TNFGlc (TG) vs siCtr_no in four biological processes: YAP/TAZ-specific and MSigDB IDs M5897, M5913 and M5830 of the interaction (siCtr_TG-siCtr_no)-(siYT_TG-siYT_no) and the sex-TG-specific interaction ((fem_siCtr_TG-fem_siCtr_no)-(fem_siYT_TG-fem_siYT_no)-(male_siCtr_TG-male_siCtr_no)-(male_siYT_TG-male_siYT_no)). X-axis is the list of all genes sorted by their P value. Y-axis is the cumulative fraction. Light blue and light red colors represent significant down– or up-regulation, respectively. (**B**) Heatmaps display selected differentially expressed genes (log2FC > 1; FDR < 0.05). (**C-D**) Integrative OMICs analysis showing boxplots of DEGs similarly (**C**) or differentially (**D**) regulated on RNA (upper panel) and protein level (lower panel). Adj. P values < 0.05 (*), <0.005 (**), <0.0005 (***), <0.0001 (****)

**Extended Data Figure 4. Quantification of microvascular network-on-chip phenotypes.** (**A**) Representative images of the microvascular network-on-chip assay at day six using male-derived HUVECs subjected to either no treatment (no) or TNF/Glc treatment (TG). Grayscale images show maximum-intensity projections of the segmented signal. Binary-colored masks represent signal segmentation, with individual objects color-coded. Intensity scale bars are shown on the grayscale images; scale bar, 100 µm. Orange rectangles indicate regions shown as 3D renderings in Fig. 6C. (**B**) Comparison of female-derived (fem, red dots) and male-derived (male, blue dots) HUVECs during microvascular network-on-chip under no treatment (no) or TG. Based on the surface area of segmented objects, data were categorized into three groups: Ruptures, disconnected cells, and small vascular connections (1,000–10,000 µm²); medium-sized vascular connections (10,000–100,000 µm²); large vascular connections (>100,000 µm²). Data represent N=14 (fem no, fem TG), N=13 (male no, male TG), analyzed from three independent experiments, P < 0,0005 (***).

**Suppl.Tab.S1.** List of taqman probes used for qRT-PCR analysis.

| target | Assay_ID |
| --- | --- |
| ANKRD1 | Hs00923599_m1 |
| CTGF<br>(CCN2) | Hs00170014_m1 |
| CXCL8 | Hs00174103_m1 |
| ESM1 | Hs00199831_m1 |
| FSP1 | Hs00243202_m1 |
| ICAM1 | Hs00164932_m1 |
| IL6 | Hs00174131_m1 |
| NOX4 | Hs01379108_g1 |
| PAI1 | Hs00167155_m1 |
| TEAD3 | Hs00243231_m1 |
| TGFBR3 | Hs00234257_m1 |
| VGLL3 | Hs01013371_m1 |
| VGLL4 | Hs00893985_m1 |
| WWTR1 | Hs00210007_m1 |
| YAP1 | Hs00902712_g1 |
| GAPDH | Hs02758991_g1 |

**Suppl.Tab.S2.** List of antibodies and dyes used for immunofluorescence.

|  | Reference | Dilution | Company |
| --- | --- | --- | --- |
| Yap | PA1-46189 | 1:100 | Thermo Fisher |
|  | sc-271134 | 1:100 | Santa Cruz Biotechnology |
| Taz | HPA007415 | 1:100 | Sigma |
| VE-Cadherin | AF938 | 1:400 | R&D sytems |
| KLF4 | HPA002926 | 1:100 | Atlas Antibodies |
| *Secondaries IF* |  |  |  |
| Donkey anti-goat Alexa Fluor 488 | A11055 | 1:250 | Thermo Fisher |
| Donkey anti-goat Alexa Fluor 647 | A21447 | 1:250 | Thermo Fisher |
| Donkey anti-rabbit Alexa Fluor 488 | A21206 | 1:400 | Thermo Fisher |
| Donkey anti-mouse Alexa Fluor 488 | A21202 | 1:400 | Thermo Fisher |
| Donkey anti-rabbit Alexa Fluor 568 | A10042 | 1:400 | Thermo Fisher |
| Donkey anti-mouse Alexa Fluor 568 | A10037 | 1:400 | Thermo Fisher |

