## Extended data figures 1- 4 for "Endothelial YAP/TAZ rewiring under cardiometabolic stress drives sex-divergent vascular remodeling in heart failure with preserved ejection fraction"

#### Extended Data Figure 1

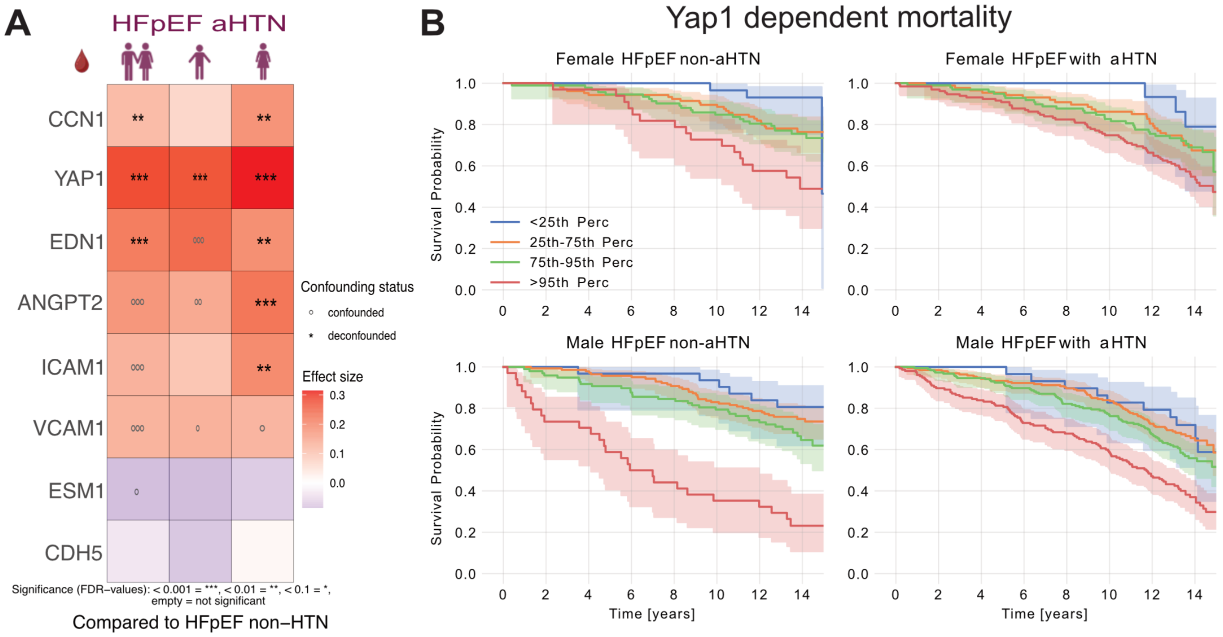

#### Extended Data Figure 2

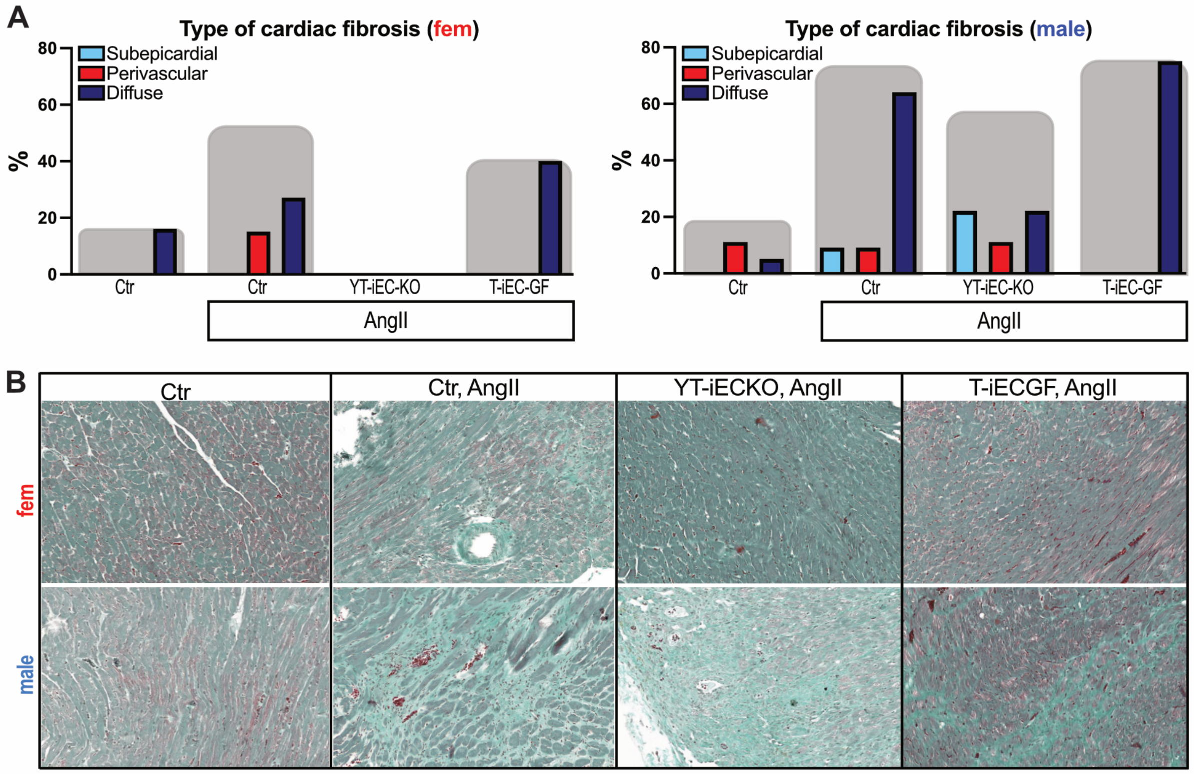

#### Extended Data Figure 3

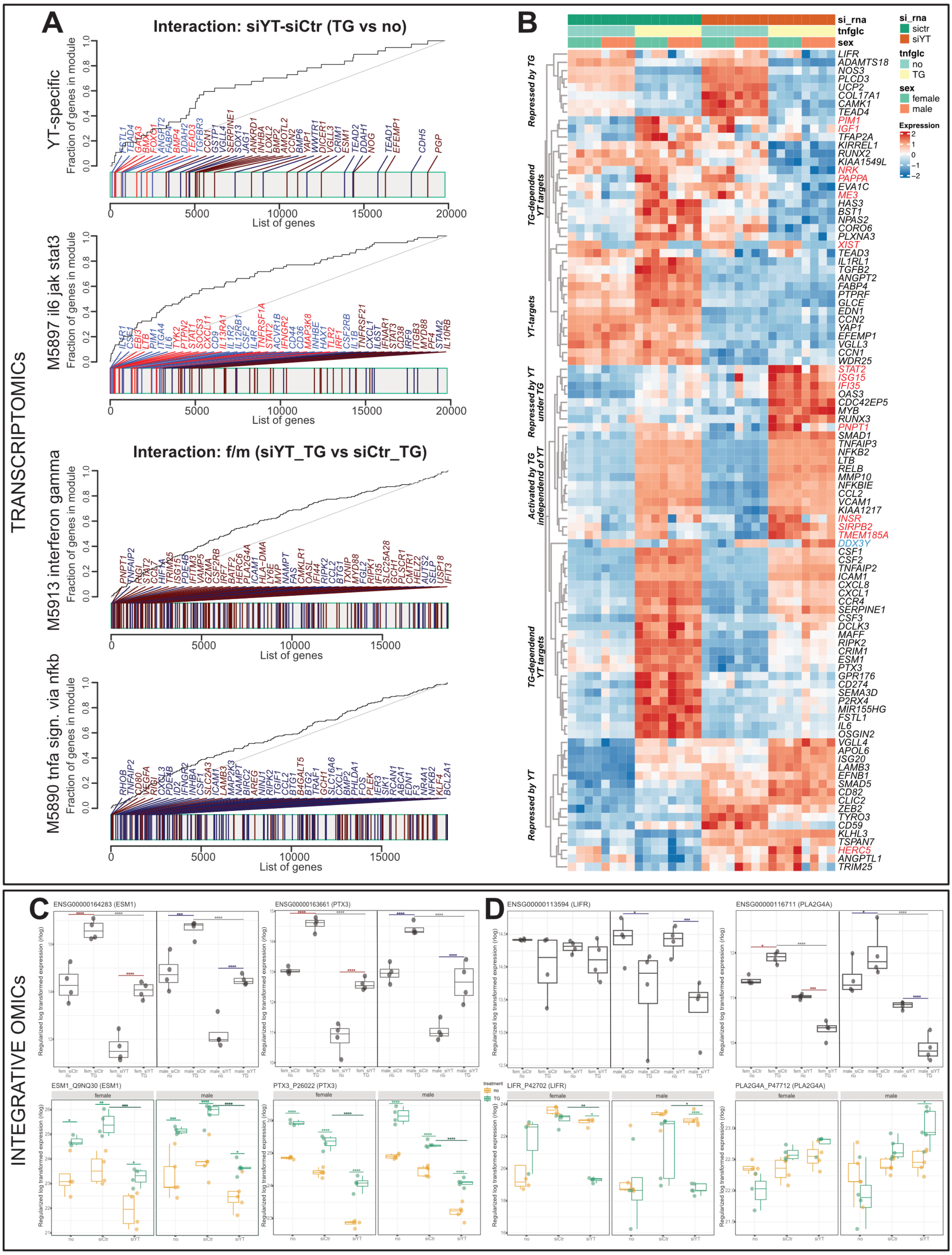

### Extended Data Figure 4

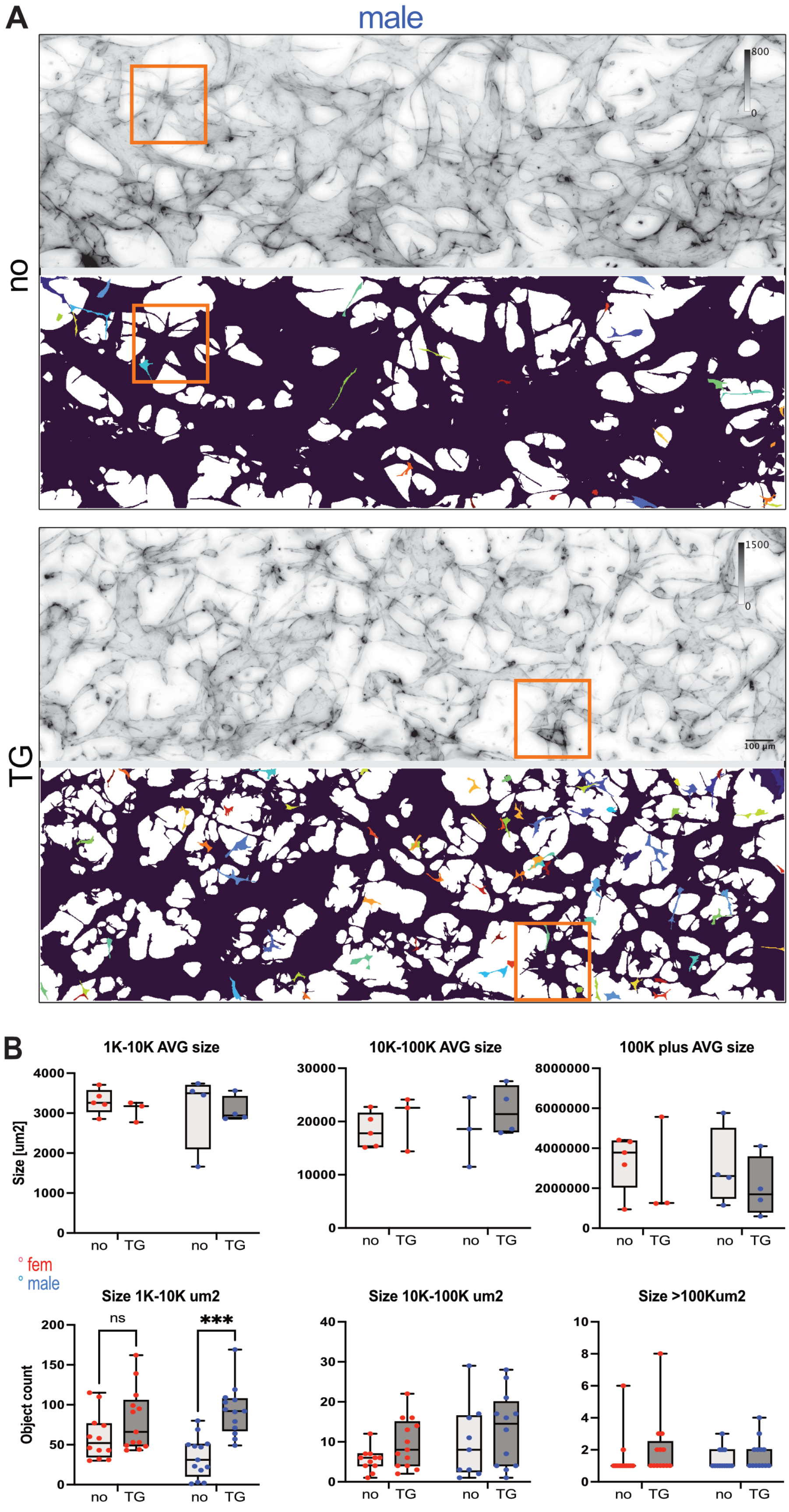
